# Pannexin 1 inhibition reduces tumorigenic properties of patient-derived glioblastoma cells through the HIPPO and Wnt signalling pathways

**DOI:** 10.64898/2026.09.03.749222

**Authors:** Danielle Johnston, Matthew Huver, Rehanna Kanji, Carlijn Van Kessel, John Kelly, Rafael Sánchez-Pupo, Brooke L. O’Donnell, Norah Defamie, Rebecca Lau, Carolina Herrera, Andrew Deweyert, Marc Mesnil, John Ronald, Matthew Hebb, Silvia Penuela

## Abstract

Glioblastoma (GBM) is the most common primary brain tumour, with a median survival of 12-18 months, highlighting a need for new treatment targets. We observed that pannexin 1 (PANX1), a channel-forming glycoprotein important in purinergic signalling, is upregulated in GBM compared to normal tissue and expressed throughout patient tumours. Western blot analysis of patient-derived GBM cell lines revealed significantly increased PANX1 expression in these primary lines compared to brain tissue and control glial cells. Bulk RNA-sequencing compared the gene expression of GBM cells devoid of PANX1 via CRISPR/Cas9 deletion (PANX1-KO) compared to controls. Gene Ontology and KEGG gene set analyses revealed PANX1-KO in GBM cells affects cell surface and cell junction components, processes, and pathways, including the HIPPO pathway, in addition to critically downregulating β-catenin mRNA and other components of the Wnt pathway. The deletion of PANX1 resulted in a disruption of the β-catenin protein and a dramatic reduction in migration and cell growth. Pharmacological inhibition of PANX1 in GBM cells with Probenecid (PBN) and Spironolactone (SPIR) demonstrated a significant reduction in live cell numbers and migration via scratch assay. Both blockers dramatically decreased F-actin filament formation, and the cellular localization of beta-catenin became more intracellular compared to controls. Xenografted GBM tumours showed a reduction in tumour cell viability by bioluminescent imaging and reduced hemorrhaging incidence when treated with PBN. These new insights support further investigation of PANX1 as a potential GBM therapeutic target and its role in multiple cancer signaling pathways that regulate this devastating disease.

## INTRODUCTION

Glioblastoma multiforme (GBM) is the most prevalent malignant central nervous system (CNS) tumour in adults, with an estimated incidence rate of 4.49 per 100,000 persons annually [1]. Classified by the World Health Organization as a grade 4 diffuse glioma, GBM is marked by rapid growth, intratumoral heterogeneity, and extensive invasion into surrounding tissue [2]. The current standard of care for GBM involves maximal safe resection of the tumour with adjuvant radiotherapy and chemotherapy, typically using the agent temozolomide (TMZ) [3]. Despite available treatment options, GBM recurrence is nearly ubiquitous among patients, and the prognosis remains poor, with a median survival of 12-18 months [4]. Thus, GBM remains an incurable, highly aggressive form of cancer, highlighting a dire need for effective therapeutics to improve patient outcomes.

Pannexins (PANX 1, 2, 3) are a family of large pore-forming glycoproteins that facilitate the passage of ions and metabolites across cell membranes [5]. PANX1 is expressed in neuronal and glial cells within the CNS, facilitating paracrine signalling [6]. Its expression levels peak during early CNS development and gradually decrease into adulthood [7, 8]. In contrast, PANX2 increases during postnatal development and remains stable throughout adulthood, while there is no evidence of PANX3 expression in the CNS [9]. PANX1 is the most well-characterized member of the pannexin family in physiological function and disease. Canonically, PANX1 facilitates the transport of ions and small molecules of less than 1 kDa across the endoplasmic reticulum (ER) and plasma membranes [5]. Activation of PANX1 is stimulated by several mechanisms, including caspase cleavage, elevated extracellular K^+^, intracellular Ca2^+^, receptor-mediated stimulation (P2X7, α1D-AR), and mechanical stimulation, and further modulated by post-translational modifications to the PANX1 channel [10–14]. At the cell surface, PANX1 participates in regulatory crosstalk with purinergic receptors through mediating cellular ATP release [13, 15]. Beyond its role at the plasma membrane, PANX1 also functions as a mechanically responsive Ca^2+^ leak channel at the ER [16]. Additionally, several PANX1 interactors have been studied, highlighting a channel-independent role of PANX1. Bhalla-Gehi *et al.* showed that disrupting actin microfilaments leads to intracellular retention of PANX1 [17]. They confirmed that the C-terminus of PANX1 directly interacts with filamentous actin (F-actin), implicating the cytoskeleton in PANX1 trafficking and stability [17]. Another group confirmed that PANX1 interacts with actin-related protein 3 (Arp3), which is involved in actin remodelling and is crucial in cell motility [18].

PANX1 dysregulation has been studied in numerous cancer types, including melanoma, breast cancer, colorectal cancer and glioma, leading to a growing interest in understanding the regulatory role PANX1 may play in tumorigenesis [19–22]. Evidence linking the expression of PANX1 and glioma was first revealed by Lai et al., wherein endogenous PANX1 expression was absent from rat C6 glioma cells; however, ectopic PANX1 expression induced a flattened morphology, reduced cell motility and decreased anchorage-dependent growth [23]. Bao and colleagues later found that ectopic PANX1 expression in C6 glioma cells unexpectedly accelerated the formation of multicellular aggregates [24]. They established that ATP release by PANX1 contributed to the activation of P2X_7_ purinergic receptors, which promoted the development of the F-actin cytoskeleton network, leading to increased aggregate formation [24]. Furthermore, PANX1 expression was detected in proliferating human glioma cells (U87-MG) and silencing PANX1 using short interfering RNA (siRNA) significantly decreased proliferation [22]. A more recent study demonstrated that overexpression of Circ-TLK1 promoted PANX1 expression via suppression of miR-17-5p in glioma, which markedly induced proliferation, invasion and migration of glioma cells [25].

To date, the role of PANX1 in Wnt/β-catenin signalling is suggested as a notable contributor to the tumor-promoting properties of PANX1 [26]. In melanoma, PANX1 directly binds and stabilizes β-catenin expression. This relationship promotes tumour growth and impairs cellular metabolism [26]. Taken together, this evidence supports the potential advantages of inhibiting PANX1 as an anti-cancer intervention.

Pharmacological inhibition of PANX1 has commonly been used to elucidate the role of PANX1 in diverse physiological contexts. Probenecid (PBN) and spironolactone (SPIR) are two established PANX1 inhibitors that cross the blood brain barrier and are FDA and Health Canada-approved drugs predominantly used for the treatment of gout and hypertension, respectively [27, 28]. PBN was shown to inhibit PANX1 channels in a non-selective manner via a binding interaction with the first extracellular loop of PANX1 [29]. Comparatively, SPIR is a specific inhibitor of PANX1 channels, predicted to interact with the C-terminus [28].

In this study, we explored the effects of inhibiting PANX1 in patient-derived GBM cells using the CRISPR/Cas9 system for a complete knockout (KO) and PBN/SPIR treatment to moderately block channel activity. It was hypothesized that targeting PANX1 would reduce its tumour-promoting effects and disrupt cellular processes that support GBM tumorigenicity. We confirmed that PANX1 is highly expressed in patient-derived GBM cells compared to non-neoplastic brain, and that inhibiting PANX1 alters cellular pathways involved in cell growth and motility. Inhibiting PANX1 dysregulated β-catenin expression along with F-actin organization. We further demonstrated that inhibiting PANX1 reduces GBM viability and hemorrhaging incidence in the *ex-vivo* chick chorioallantoic membrane (chick-CAM) assay. These new insights support further investigation into PANX1 as a potential target for novel GBM therapeutics.

## METHODS

### Primary glioblastoma cells and culture conditions

Patient-derived glioblastoma and control cells from brain biopsies [30] were collected by the Hebb Lab (Lawson Health Research Institute) from consented patients under REB 17290 as described previously [31] Immunoblotting analysis of the obtained samples confirmed that the patient sample GBM17 cells appreciably expressed PANX1. GBM17 cells were used as representative glioblastoma cells for this study. Cells were cultured in Dulbecco’s Modified Eagle Medium 1X (DMEM 1X, Thermo Fisher Scientific) containing 4.5% g/L D-glucose, L-glutamine, and 110 mg sodium pyruvate, supplemented with 10% fetal bovine serum (FBS, Wisent), 100 units/mL penicillin, and 0.1 mg/mL streptomycin, and incubated at 37°C and 5% CO2. Trypsin (0.25%, 2.21 mM EDTA; Wisent) was used to dissociate cells from culture plates. Commercial glioblastoma cell lines U87 and LN229 were obtained from ATCC (U87 #CRL- HTB-14; LN229 #CRL-2611) and U251 was obtained from Millipore Sigma (#09063001-1VL). All commercial lines were cultured in DMEM as described above.

### CRISPR/Cas9-generated PANX1 knockout

The CRISPR/Cas9 D10A system [32] was used to generate patient-derived GBM17 *PANX1*- KO cells using plasmids gifted by Feng Zhang (Broad Institute, MIT) (Addgene plasmid #48140 and #62987). GBM17 cells were cultured in 6-well plates (Corning, #3506), then transfected with 1 µg pSpCas9n(BB)-2A-Puro (PX462) V2.0 and 1 µg pSpCas9n(BB)-2A-GFP (PX451) containing guide RNA with the sequences: GTTCTCGGATTTCTTGCTGA to target *PANX1* exon 1 and CTCCGTGCCAGTTGAGCGA as a scramble control Guide RNA sequences were designed using tools.genome-engineering.org. Cell selection was completed using 1 µg/mL puromycin for 72 hours, 24 hours after transfection, then single colonies were screened with immunoblotting probing for PANX1 expression.

### Illumina next generation sequencing

RNA was isolated from 3 PANX1-KO GBM17 clones and 3 control GBM17 clones using an RNeasy mini kit (Qiagen, #74104) and Bulk RNA-sequencing (RNA-seq) was completed at the London Regional Genomics Centre (Robarts Research Institute, London, Ontario, Canada; http://www.lrgc.ca).

A NanoDrop 1000 (Thermo Fisher Scientific) was used to quantify total RNA from samples. The quality of the samples was assessed using Agilent 2100 Bioanalyzer (Agilent Technologies Inc.) and the RNA 6000 Nano kit (Caliper Life Sciences) with 1 µL (50-500 ng/µL) of sample per GBM clone. Library preparation for sequencing, including rRNA reduction was completed using a Vazyme VAHTS V8 RNA-seq Library Prep Kit for Illumina (Vazyme, #NR-60502).

cDNA from each sample was prepared by rRNA depletion and fragmentation, followed by indexing, before amplification via PCR. The generated libraries were then equimolar pooled into one library, evaluated for size distribution, then quantified using an Agilent High Sensitivity DNA Bioanalyzer chip and a Qubit 2.0 Fluorimeter (Thermo Fisher Scientific), respectively. An Illumina NextSeq500 (Illumina Inc., San Diego, CA) with a MidOutput v2 kit (**#**20024904, 150 cycles, 2 x 76 bp paired end) was used to sequence the library.

Partek Flow Version 11.0.24.022 (St. Louis, MO) was used for subsequent analysis of Fastq Files, including alignment of the data to the *Homo sapiens* genome with STAR 2.7.3a [33] and annotation to hg38 Ensembl Transcripts release 109 [34] using the quantify to annotation model (Partek E/M). The standard pre- and post-alignment QA/QC provided in Partek Flow was completed. Detected genes with a total read count less than 10 were filtered out before applying DESeq2 [35] for normalization and analysis of the counts data to generate p-value and fold change values. A filtered list of detected genes was generated in Partek Flow to include features that exhibited a fold change greater than absolute 1.5 and a false discovery rate (FDR) step-up of less than 0.05, where FDR step-up is representative of the adjusted p-value. Gene Ontology (GO) [36] and Kyoto Encyclopedia of Genes and Genomes (KEGG) [37] pathway enrichment analyses were performed in R using filtered lists of differentially expressed genes (DEGs). Enrichment was assessed using R-based functional annotation tools [38], and the top enriched biological processes, cellular components, molecular functions, and KEGG pathways were visualized as bubble plots according to adjusted p-value and gene count. The principal component analysis (PCA) was completed using Partek Flow. evaluating DEGs with an FDR step-up of less than 0.05 and a fold change greater than absolute 1.5.

#### GSEA

Gene Set Enrichment Analysis (GSEA) [39] was conducted using the normalized RNA-seq data collected from DESeq2 analysis in Partek Flow, evaluating *a priori* defined Molecular Signature Database (MSigDB) human hallmark gene set collection [40]. Normalized gene counts were rounded to the nearest whole number for analysis in GSEA software. Corresponding enrichment plots, heatmaps and leading-edge analysis were generated using GSEA software.

*Monolayer (2D) cell culture assays and drug Treatment*PBN (Thermo Fisher Scientific, #P36400) dissolved in water or SPIR (Selleckchem, S4054) dissolved in ethanol at a working concentration of 1mM and 20µM, respectively, were used as treatments to inhibit PANX1 channels in vitro. For monolayer cell culture assays investigating PANX1 channel inhibitors, GBM17 cells were plated in 24-well plates for live-cell imaging and 6-well plates (Corning, #3506) for immunofluorescence analysis and protein and RNA extraction, at seeding densities of 20,000 and 80,000 cells per well, respectively. Cells were treated 3 times over 5 days with PBN, SPIR, water, or ethanol, beginning 24 hours following cell plating, by replacing the used culture medium with fresh culture medium mixed with the respective blocker or vehicle control.

*Live cell imaging*Twenty-four-well and 96-well plates were incubated in an Incucyte® Live-Cell Analysis S3 System (Sartorius, #4763) over the course of treatment. Phase contrast images of each well were acquired with an embedded brightfield microscope within the Incucyte® at a 4x and 10x objective lens magnification. Post-treatment analysis using the Incucyte® software included cell count for monolayer cell culture.

### Protein extraction and immunoblotting

Protein lysates were extracted from monolayer cell cultures with RIPA buffer (50 mM Tris-HCl pH 8.0, 150 mM NaCl 1% NP-40 [Igepal], 0.5% Sodium Deoxycholate, 0.1% SDS) containing 100 mM Sodium Fluoride and 100 mM Sodium Orthovanadate dissolved in ddH2O, and half a tablet of complete-mini EDTA-free Roche Tablet. Protein concentration was quantified via a Bicinchoninic acid (BCA) Protein Assay (Thermo Fisher Scientific, #23225), and samples were mixed with SDS sample buffer (4X) prior to protein electrophoresis. Protein lysates (50 µg) and a Precision Plus Protein™ Prestained Protein Standard (Bio-Rad, #1610375) were separated using a 10% SDS-PAGE gel, which was then transferred onto a nitrocellulose membrane using an iBlot transfer stack and apparatus with program P3 (Invitrogen, #IB301002). Membranes were blocked using 3% bovine serum albumin (BSA) mixed in phosphate-buffered saline (PBS) 1X and then incubated with anti-human PANX1 (1:1000, PANX1 CT-412 [41]) and anti-human GAPDH (1:5000 dilution, MilliporeSigma #MAB374) primary antibodies, followed by Goat anti-Rabbit 800 (1:10,000 dilution, LI-COR #926-32211) and Goat anti-Mouse 680 (1:10,000 dilution, LI-COR #926-68070) secondary antibodies, all diluted in PBS with 0.05% Tween20. GADPH was used as a loading control. Western blots were imaged using a Li-COR Odyssey Classic infrared imaging system and quantified using Odyssey Application Software (Version 3.0, LI-COR). Blots were re-incubated proceeding the first scan with anti-human β-catenin (1:1000, Cell Signaling Technology #9562) or anti-human β-actin (1:2000, Millipore-Sigma #A2228) primary antibody and corresponding above secondary antibodies, then re-imaged and quantified.

### Quantitative reverse transcription PCR (RT-qPCR)

Cell samples for RNA extraction were collected from 6-well (monolayer cell culture) and 96- well (tumour spheroids) plates post-treatment and centrifuged to form cell pellets, then stored at - 80°C until further use. RNeasy® Plus Mini Kit (Qiagen, #74904) was used to purify total RNA from each cell sample. β-mercaptoethanol was added to the Buffer RLT plus before use, as indicated in the Quick-Start Protocol. RNA lysis purity was assessed using a microplate spectrophotometer (BioTek). RNA samples were prepared for reverse transcription using a High-Capacity cDNA Reverse Transcriptase Kit (Thermo Fisher Scientific, #4368814) according to the manufacture’s protocol. A T100 Thermal Cycler (Bio-Rad) was used for cDNA synthesis. The following DNA primers (Invitrogen) were obtained: hPANX1l and hPANX1u, hB-cat 1F and hB-cat 1R, IL-8 F and IL-8 R, human GAPDH R and human GAPDH F, Actb - F and Actb -R. GADPH was used as an expression control. RT-qPCR was conducted with SsoAdvanced Universal SYBR® Green Supermix (Bio-Rad, #1725274) mixed with each cDNA sample and respective primer mix. RT-qPCR samples were run in triplicates, using a CFX Connect Real-Time PCR Detection System (Bio-Rad). Fold change for each gene of interest was calculated using the ΔΔCt method.

### Immunofluorescence microscopy

Monolayer cell cultures were grown on coverslips and treated in parallel with the live-cell imaged plates over 5 days. Cells were fixed in 80% methanol/20% acetone at 4°C for 15 minutes or 4% paraformaldehyde for 30 minutes at room temperature, with 0.1% TritonX100 was added to the blocking and antibody buffers throughout the remaining procedure. All samples were blocked with 2% BSA in 1X PBS and incubated with anti-human PANX1 (1:500, PANX1 CT- 412), and phalloidin-568 (1:20, Thermo Fisher Scientific #A12380) or anti-human β-catenin (1:100, BD Transduction Laboratories #610154) primary antibodies. Hoechst 33342 (1:1000, Thermo Fisher Scientific #62249) was used to stain nuclei, Alexa Fluor 647 Goat anti-Rabbit (1:400, Thermo Fisher Scientific #A-21244) and Alexa Fluor 488 Goat anti-Mouse (1:700, Thermo Fisher Scientific #A-32723) were used as secondary antibodies for PANX1 and β- catenin, respectively. Coverslips were mounted on glass with ProLong Gold (Thermo Fisher Scientific, #P36930) prior to imaging. Immunofluorescence images were obtained using a Zeiss LSM 800 laser scanning confocal microscope at 63x (Carl Zeiss).

### Immunohistochemistry

GBM tumours and patient-matched non-neoplastic brain tissues were fixed in 10% formalin overnight at 4 degrees. All tissues were sent to Robarts Research Institute for processing and embedding. Serial sections of 5 µm were used for hematoxylin & eosin (H&E) and 3,3’- diaminobenzidine (DAB) staining. For DAB staining, tissue slides were incubated at 40°C overnight, followed by xylene deparaffinization and decreasing ethanol (EtOH) concentrations for rehydration. Endogenous peroxidases and alkaline phosphatases were quenched using BIOXALL Endogenous Blocking Solution (Vector Laboratories #SP-6000) for 10 minutes. Sections were washed with 1X PBS for 5 minutes, and antigen retrieval was performed with Antigen Unmasking Solution Citrate-based pH 6.0 (Vector Laboratories #H-3300) in a decloaking chamber at 112.5°C for 1.5 minutes and 90°C for 10 seconds. Sections were washed with 1X PBS for 5 minutes and then blocked with 2.5% Normal Horse Serum for 30 minutes at room temperature (Vector Laboratories #S-2012). Blocking solution was left on secondary only control slides while the remaining slides were incubated with 1:250 PANX1 CT- 412 diluted in 2.5% Normal Horse Serum overnight at 4°C. All slides were washed twice in 1X PBS for 5 minutes and incubated with ImmPRESS HRP Horse Anti-Rabbit IgG Polymer Reagent (Vector Laboratories MP-7401) for 30 mins at room temperature, followed by two washes in 1X PBS for 5 minutes. DAB substrate (Vector Laboratories SK-4100, using reagents 1–3) was prepared in distilled water and incubated on sections for 3 minutes. Slides were rinsed with distilled water followed by a wash in tap water. Slides were counterstained with Harris hematoxylin (Leica 3801561, Wetzlar, Germany) for 1 minute, followed by a rinse in tap water. Slides were dipped thrice in 0.5% acid alcohol, rinsed in tap water, dipped three times in 2% ammonium alcohol and rinsed in tap water again. Slides were cleared using increasing concentrations of alcohol and xylene followed by mounting using Fisher Chemical Permount Mounting Medium (Thermo Fisher Scientific SP15) and microscope cover glass. The slides were scanned using an Aperio Glass Slide Scanner.

### MTT cell viability assay and Cytotox growth curves

A 3-(4,5-dimethylthiazol-2-yl)-)-2,5-diphenyltetrazolium bromide (MTT) assay (Abcam # ab211091) was used to confirm treatments of PBN (1 mM) and SPIR (20 µM) did not affect GBM17 cell viability. A total count of 50,000 cells were plated in 24-well plates. Forty-eight hours after treatment, MTT was diluted 1:40 in culture medium and incubated with the cells for 1 hour at 37°C with 5% CO_2_. For both experiments, culture medium was aspirated, and 500 µL of DMSO was added to each well and shaken for 5 minutes at room temperature to dissolve the formazan products. In a 96-well plate, triplicates of each condition were diluted 1:2 in DMSO with 100 µL of DMSO in triplicate serving as blanks. OD570 was measured using a VICTOR3 Multilabel Counter plate reader (PerkinElmer). Triplicate values were averaged for each experiment. Cytotox growth curves were completed by plating GBM17 cells at 20,000 cells per well in a 24-well plate (Greiner Bio-One, #662165). PBN (1mM) dissolved in H_2_O, SPIR (20µM) dissolved in ethanol, TMZ (50µM) dissolved in dimethyl sulfoxide (DMSO) or vehicle controls, H_2_O, ethanol and DMSO, were administered to cells three times over five days, beginning 24 hours after plating. Cells were treated with Cytotox Red Dye (Sartorius, #4632) along with the previous treatments. Nine images were captured per well every 4 hours. The Incucyte S3 System software was used to obtain red counts or phase counts for cytotoxicity curves and growth curves, respectively.

### Fluorescent-gelatin degradation assay on insert

To assess the ability of cells to form invadopodia and degrade matrix, LN229 cells were plated on inserts (10^4^ cells/mL) coated with 0.2% fluorescent FG-gelatin (Molecular Probes, #G13186) in 24-well plates and maintained in an incubator (37°C, 5% CO_2_) for 8 hours. The cells were fixed in paraformaldehyde (4%) for 20 min at room temperature. After incubation in a blocking solution (2% bovine serum albumin, 1% TritonX100 in PBS), the cells were incubated with primary antibodies (anti-actin 1:250 and anti-PANX1 CT-412 1:500) overnight at 4°C.

Mouse Alexa Fluor® 555- and Rabbit Alexa Fluor® 647-conjugated antibodies (1:250, Invitrogen) were then applied on the preparations for 1 hour. Coverslips or inserts were mounted afterwards with Mowiol (Millipore Sigma, #81382) prior to observation with confocal microscopy. As a first step, the presence of invadopodia was checked at the level of digested areas of green-fluorescent gelatin (FG-gelatin; 488 nm) which appear as black spots. The second step was to identify, at the level of the black spots, invadopodia through their molecular components (actin), which were detected by red labeling (545 nm). Thereafter, the third step was the detection of PANX1 by far red labeling (633 nm). Confocal images were obtained using an Olympus IX81 laser scanning confocal inverted microscope with 40X UAPO ID/340UV NA 135 oil or 60X O.N. 1.4 PLAPO oil objectives. Images were processed with FluoView software.

### Xenograft Tumor Growth in the Chick Chorioallantoic Membrane Assay (Chick-CAM Assay)

Chick-CAM assays were performed as described previously [19, 42]. Briefly, fertilized chicken eggs (McKinley Hatchery, St. Mary’s, ON, Canada) were incubated in a rotating incubator for 3 days. Embryos were transferred into weigh boats and incubated for 7 more days. GBM17 cells were transduced with lentivirus containing tdTomato-firefly luciferase. At day 10, 1 x 10^6^ cells in serum-free media were combined with Matrigel (Corning #356237) (1:1) and implanted onto a branching vessel of the CAM. Tumors were treated daily with a topical application of 1mM PBN or vehicle. At day 17, luciferin was applied to tumors, and they were imaged with bioluminescent imaging (BLI) on an IVIS Lumina XRMS to assess for cell, viability. For histological analysis, tumors were excised on day 18 of the experiment, retaining the surrounding CAM microenvironment, fixed in 10% formalin and embedded in paraffin as previously described [19]. The number of chicks that developed hemorrhaging at day 18 endpoint was counted and used to calculate the incidence of hemorrhaging as well as the odds ratio of developing hemorrhaging.

### Statistical analysis

Statistical analyses of collected data were performed using GraphPad Prism (Version 9.3). Data collected from live cell imaging were evaluated using a two-way ANOVA analysis, followed by Tukey’s multiple comparisons test. Quantified protein expression levels from immunoblotting were compared using a one-way ANOVA, followed by Tukey’s multiple comparisons test. Gene mRNA expression levels were evaluated using a paired t-test for each vehicle control versus treatment, per gene. A Fisher’s exact test was used to analyze hemorrhaging incidence in the chick-CAM assay. Significance is representative of *P<0.05, and **P<0.01. Mean values were calculated from at least three biological replicates ± standard deviation.

## RESULTS

### PANX1 is highly expressed in glioblastoma

We used GEPIA (Gene Expression Profiling Interactive Analysis, http://gepia.cancer-pku.cn) to examine transcript expression of *PANX1* in glioblastoma patient tumors. *PANX1* transcript was significantly upregulated in GBM tumors compared to normal brain (Fig 1A). Since GBM is a highly heterogeneous tumor consisting of many cell populations and distinct tumor regions, we explored PANX1 expression in the GBM microenvironment by staining serial sections of GBM tumor fragments (Fig 1B). PANX1 was found to be expressed around vasculature, surrounding areas of necrosis and throughout hypercellular regions of GBM tumors (Fig 1C).

**Figure 1.**
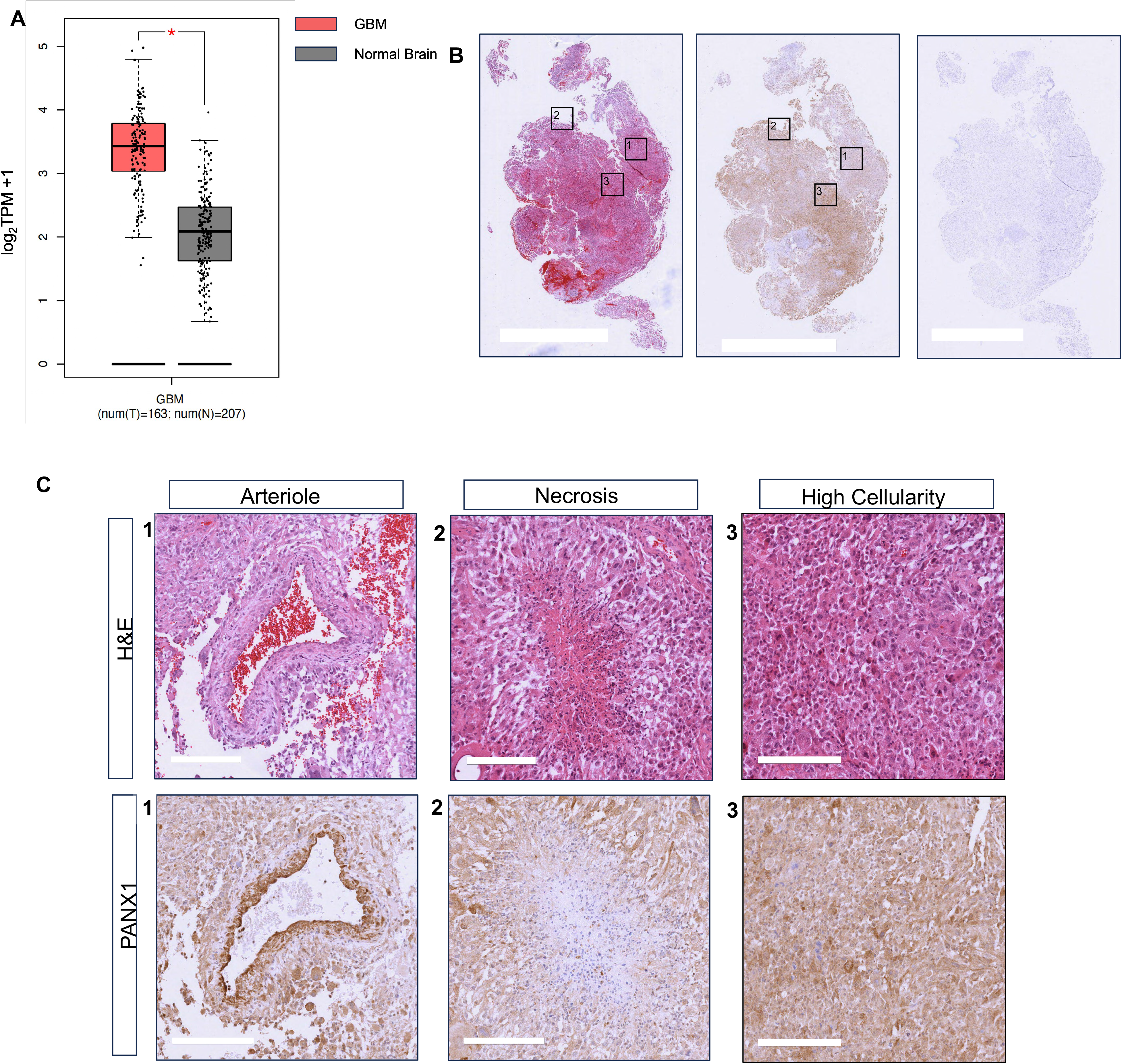
PANX1 is highly expressed in glioblastoma. **A)** Patient RNA sequencing data from GEPIA (Gene Expression Profiling Interactive Analysis). PANX1 transcript expression was significantly higher in tumor samples (N=163) compared to non-neoplastic brain tissue (N=207). **B**) Serial sections of a GBM tumour fragment stained with hematoxylin & eosin (H&E), PANX1 immunohistochemistry (IHC), and a secondary-only control. Scale bar = 5mm . **C**) 10X magnification images of the H&E and PANX1 IHC stained tumor, highlighting PANX1 localization around a blood vessel, around a necrotic area and showing diffuse staining in an area of highly cellular area. Scale bar = 200 um

We obtained various commercially available GBM cell lines, patient derived GBM cells from tumor specimens, and primary control glial cells from non-neoplastic brain specimens. Protein lysates from these cells were ran alongside normal human brain tissue, and we observed high expression of PANX1 protein compared to normal brain and control glial cells (Fig 2A). To visualize the localization of PANX1 in GBM cells, immunofluorescence was performed and demonstrated a mostly diffuse, intracellular pattern of PANX1 protein, with small amounts visible at the cell membrane in various GBM cell lines (Fig 2B).

**Figure 2.**
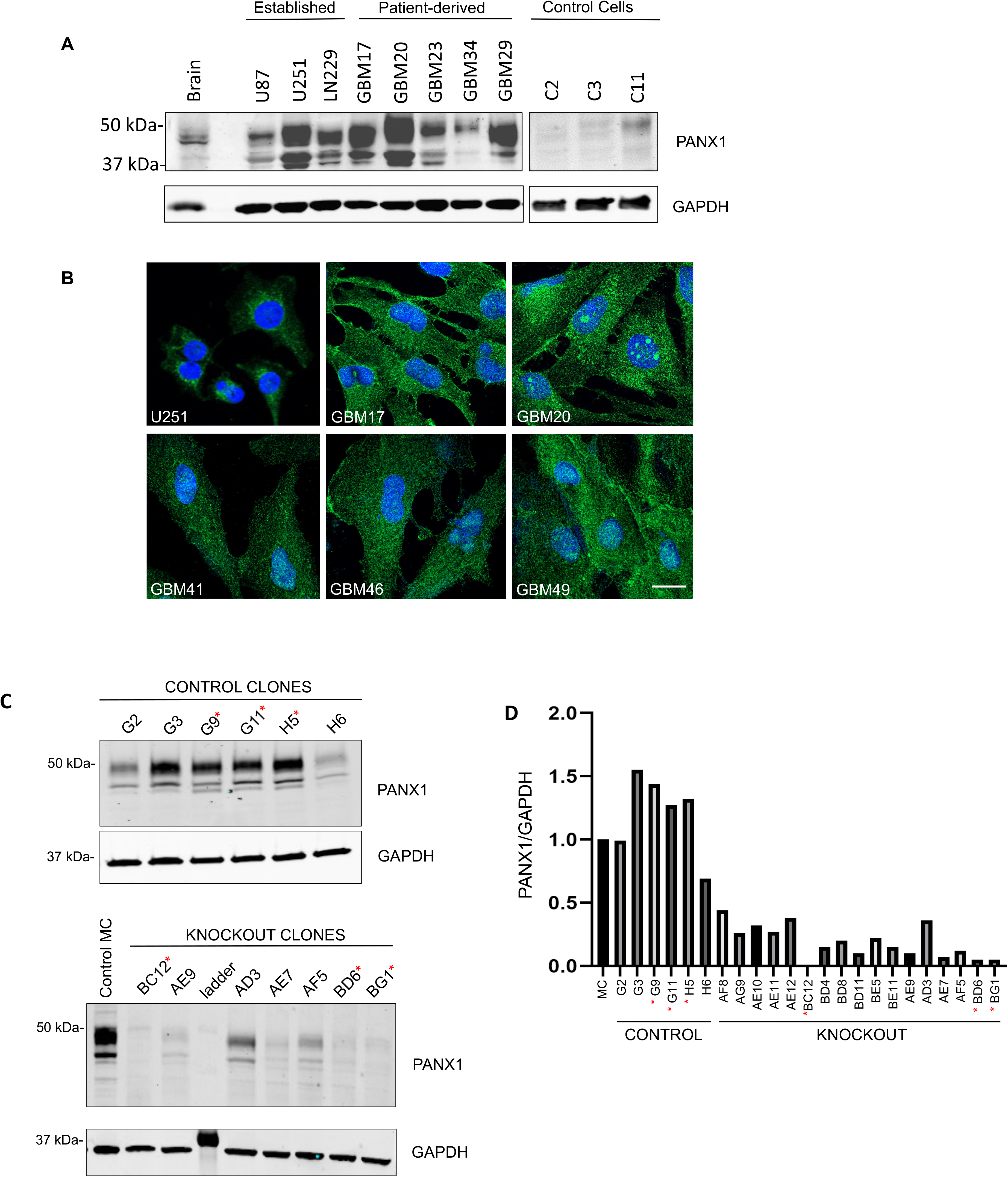
PANX1 is highly expressed in patient-derived GBM cells, PANX1 KO CRISPR clones generated for RNA-sequencing. **A)** Surgical specimens from GBM patients were used to generate primary patient-derived cell lines. A Western blot shows PANX1 protein is expressed highly in most commercial GBM cell lines and patient-derived GBM cell lines, with minimal detection in control primary glial cells. **B)** Immunofluorescence of various GBM cells (commercial and patient-derived) show majority intracellular PANX1 expression, with a small amount visible at the cell membrane (DAPI – blue, hPANX1 – green, scale bar = 20 uM).**C)** Western blot of several PANX1 KO and control clones, compared to GBM17 control mass culture (MC). Clones were screened for RNASequencing analysis. **D)** Quantification of PANX1 signal in control and KO clones. (3 KO and 3 control clones were selected).

### Differential gene expression analysis of PANX1-KO GBM cells revealed extensive gene expression changes

To elucidate the role of *PANX1* in GBM, we used CRISPR/Cas9 gene editing to knock out *PANX1* from patient-derived cell line GBM17. Single cell selection was performed on control and knockout cells, and derived cell lines were analyzed via Western blot for PANX1 levels (Fig 2C). Three control (G9, G11, H5) and 3 KO (BC12, BD6, BG1) clones were selected based on highest and lowest PANX1 protein level quantification, respectively, and RNA was purified from these cell lines for bulk RNA-sequencing. A principal component analysis (PCA) indicated that PANX1-KO GBM cell clones exhibited a similar gene expression profile phenotype between replicates and an appreciable difference compared to the control clones, forming two non-overlapping clusters visualized on the PCA plot (Suppl. Fig S2B). An MA plot with log fold change (LFC) shrinkage was generated to visualize that the normalized dataset was distributed as expected (Suppl. Fig S2C).

Evaluating the bulk RNA-seq data collected, we detected a total of 19,119 annotated genes following DESeq2 normalization. A fraction of these genes exhibited significantly altered gene expression in GBM PANX1-KO clones compared to control clones, comprised of 2,327 genes that exhibited a false discovery rate (FDR) step-up (adjusted p-value) less than 0.05 and a fold change greater than absolute 1.5. PANX1-KO GBM17 cells had 1,098 significantly upregulated DEGs and 1,229 significantly downregulated DEGs (Suppl. Fig S2A and Suppl. Fig S4). We created a hierarchical clustering heatmap to illustrate the differential expression of the top 12 DEGs and a table including the function of these proteins (Fig 3).The top 12 DEGs were *NRG1* (neuregulin 1), *ADGRL2* (adhesion G protein-coupled receptor L2), *RAP1GAP2* (RAP1 GTPase activating protein 2), *VWA1* (von Willebrand factor A domain containing 1), *TNFRSF1B* (TNF receptor superfamily member 1B), *ANGPT2* (angiopoietin 2), *PLAT* (tissue-type plasminogen activator), *SEL1L3* (SEL1L family member 3), *TIMP3* (TIMP metallopeptidase inhibitor 3), *ABI3BP* (ABI family 3 binding protein), *CREG1* (cellular repressor of E1A stimulated genes 1), and *MXRA8* (matrix remodeling associated 8). RT-qPCR of NRG1, TNFRSF1B, RAP1GAP2, ADGRL2, ABI3BP in control and PANX1-KO GBM17 cells validated the fold changes observed for each of these top DEGs identified in the RNA-seq analysis. (Suppl. Fig S2D).

**Figure 3.**
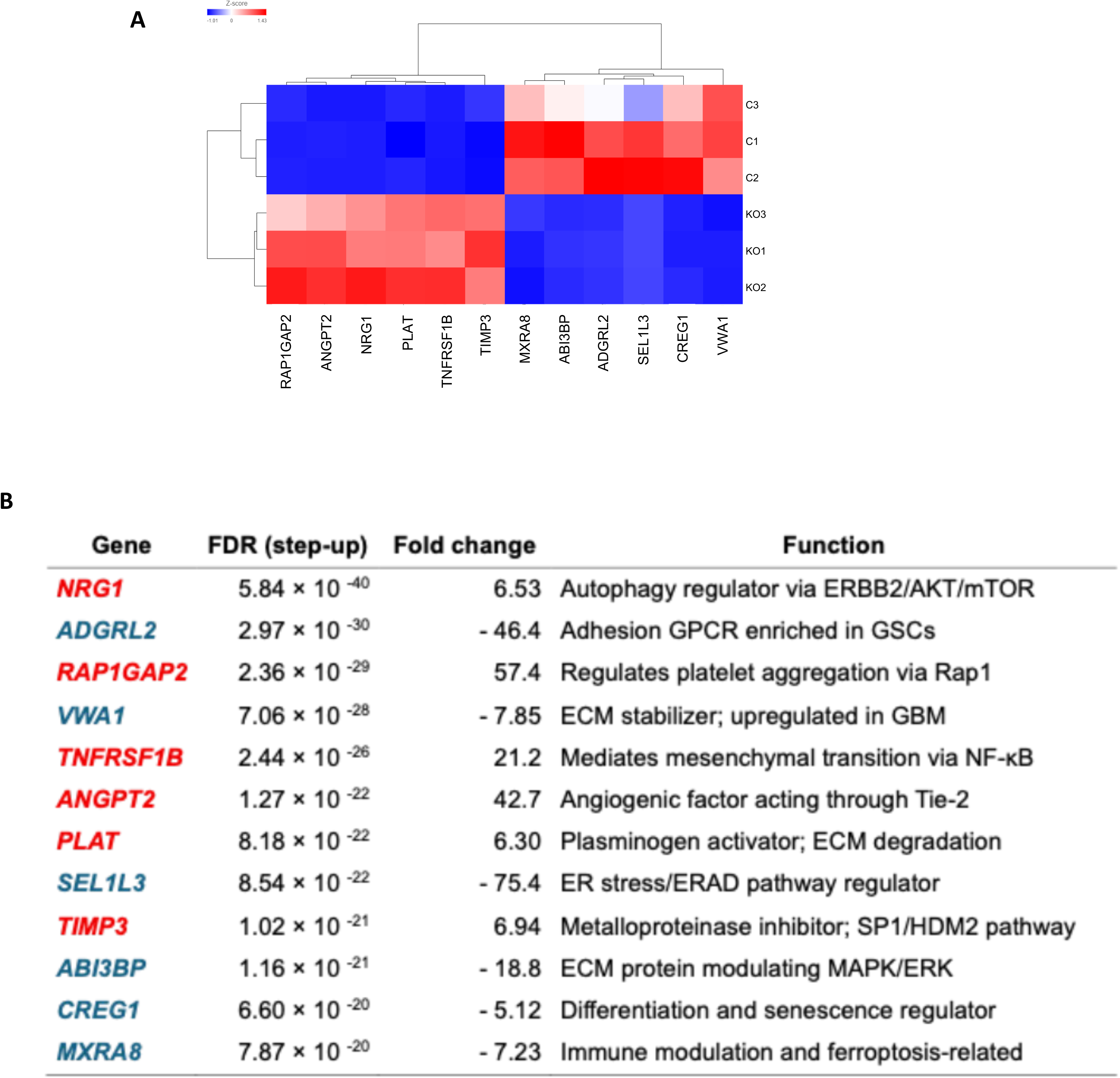
Differential gene expression analysis of PANX1 knockout GBM cells revealed extensive gene expression changes. **A)** Hierarchical clustering heatmap – Top 12 DEGs in PANX1-KO versus control GBM clones, ranked by FDR step-up. **B)** Top Differentially Expressed Genes in GBM17 PANX1-Knockout Cells Compared with Controls. Genes are ranked by FDR-adjusted significance. Fold change reflects KO relative to scrambled control (blue = downregulated, red = upregulated). Functions were summarized from the literature.

### PANX1-KO in GBM cells is associated with signaling and receptor interaction KEGG pathways

To determine the cellular pathways predominantly affected by PANX1-KO in GBM cells, we performed KEGG pathway analysis, and GSEA analysis (Suppl. Fig. S3) on our RNA-seq data, evaluating DEGs with an FDR step-up of less than 0.05 and a fold change greater than absolute 1.5. The KEGG pathway analysis revealed 4 significantly enriched pathways including ECM receptor interaction (FDR step-up = 2.34×10^-4^), PI3K-Akt signaling pathway (FDR step-up = 3.22×10^-4^), Coronavirus disease – COVID-19 (FDR step-up = 3.22×10^-4^), and the focal adhesion pathway (FDR step-up = 3.80×10^-4^). Observing the top 20 enriched pathways ordered by FDR step-up (Figure 4A), we found that multiple pathways associated with GBM tumorigenicity were altered by PANX1 KO, including PI3K-Akt signaling and MAPK signaling pathways. We noted several moderately enriched pathways that appear to coincide with the function of PANX1 at the cell surface, such as ECM-receptor interaction, neuroactive ligand-receptor interaction, and cell adhesion molecules. We detected the enrichment of viral pathology pathways, including coronavirus disease – COVID-19, Influenza A, and Human papillomavirus infection, which corresponds to previous studies that propose PANX1 may facilitate viral pathogenesis [43], [44].

**Figure 4.**
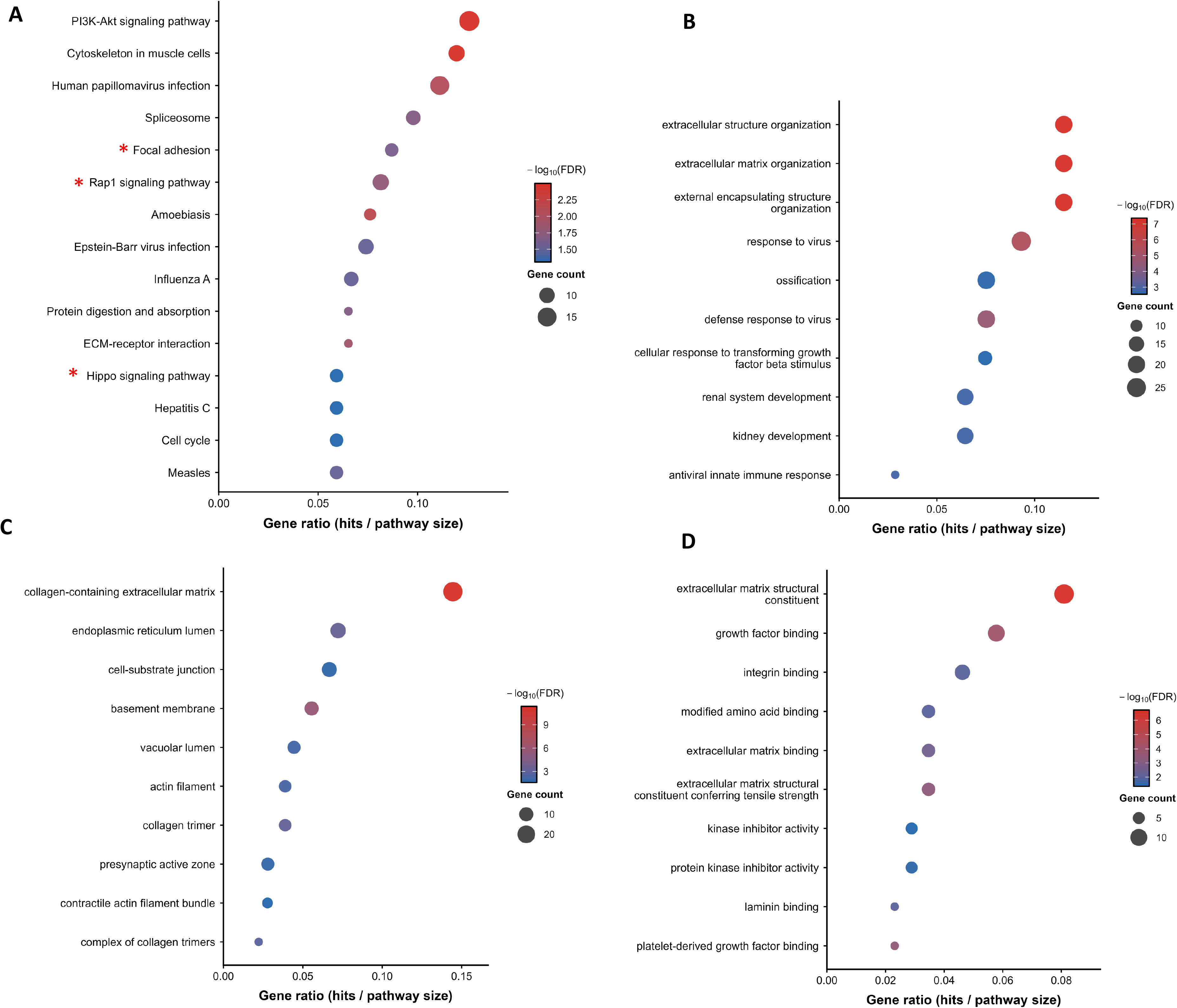
KEGG pathway analysis and Gene Ontology (GO) analysis using R. **A)** KEGG pathway enrichment - dot plot showing the top enriched KEGG pathways among differentially expressed genes (DEGs) between knockout (KO) and scrambled control (SC) samples. Dot size indicates the percentage of DEGs annotated to each pathway, and dot colour encodes the total number of DEGs per pathway. Stars indicate key pathways where β-catenin and actin are involved. **B)** GO Biological Process (BP) enrichment. **C)** GO Cellular Component (CC) enrichment. **D)** GO Molecular Function (MF) enrichment.

We found actin and β-catenin associated pathways present across multiple of the top 20 enriched pathways, including focal adhesion, Rap1 signalling, and Hippo signalling. Bhalla-Gehi *et al.* [17] and Sayedyahossein *et al*. [26] from our group, have previously determined that actin and β- catenin directly interact with the carboxyl-terminus segment of PANX1, highlighting a potential mechanism by which PANX1 ablation in GBM cells may impact these pathways. PANX1 KO in GBM cells altered the expression of genes across the entire pathway, including the significant downregulation of actin and β-catenin mRNA expression (Suppl. Fig S1). Our RNA-seq data showed β-catenin (*CTNNB1*) decreased by -1.61-fold (FDR step-up = 1.51×10^-2^) in PANX1-KO versus control GBM clones. The focal adhesion KEGG pathway map actin component critically refers to cytoplasmic actin isoforms, actin beta (*ACTB*) and actin gamma 1 (*ACTG1*) genes. Interestingly, we found that actin alpha 2 (*ACTA2*) and *ACTG1* exhibit decreased fold changes of -4.02 (FDR step-up = 2.11×10^-2^) and -1.65 (FDR step-up = 1.77×10^-2^), respectively. We detected decreased *ACTB* expression, with a fold change of -1.44 (FDR step-up of 1.81×10^-1^) showing a corresponding downward trend to *ACTA2* and *ACTG1* expression despite falling outside of the fold change and adjusted p-value cutoffs. This data highlights the protein and ECM interactions with Pannexin 1 at the cell membrane.

### Gene Ontology analysis of PANX1-KO GBM cells demonstrates predominantly plasma membrane and ECM interface interactions

To provide insight into the localization and function of the DEGs detected using RNA-seq, we completed a Gene Ontology (GO) analysis (Fig 4B-D). We conducted our analysis to include only the genes that exhibited a fold change of greater than absolute 1.5 and an FDR step-up of less than 0.05. The biological process aspect highlighted GO terms involved in intercellular interactions, indicating genes altered by PANX1 deletion are largely involved in cell adhesion and migration processes (FDR step-up ≤ 7.87×10^-9^). Enriched molecular function GO terms focused predominantly on signaling receptor activity (FDR step-up ≤ 1.56×10^-3^). The top enriched GO terms for cellular component highlighted several extracellular components, and cell and anchoring junctions (FDR step-up ≤ 3.41×10^-4^). Altogether, the GO report data suggest that PANX1-KO in patient-derived GBM cells predominantly altered the expression of genes associated with the modulation of signal receptor and cytoskeletal interaction activity at the plasma membrane and extracellular matrix interface.

### **β**-catenin protein is disrupted in KO clones and PANX1 drug inhibition reduces cell growth and migration in GBM17 patient-derived cells in 2D

The decrease in β-catenin transcript was confirmed with RT-qPCR (Fig 5A), and a disruption in the β-catenin protein was seen in the KO or KD (partial deletion/knockdown clones) via Western blot (Fig 5B). To determine how this may affect growth and motility, a scratch wound assay showed significantly decreased wound closure in the KO clone compared to control (Fig 5D). A growth curve was performed and showed a significant reduction in KO clone cell numbers compared to control (Fig 5C). However, these cells plateaued at day 4 and likely became senescent, preventing further studies. Established PANX1 blockers PBN and SPIR were used to verify if they had a similar effect as the KO on growth and motility, and both showed a significant decrease in the number of live cells and the wound closure compared to controls (although not as severe as the KO) (Fig 5 E-F).

**Figure 5.**
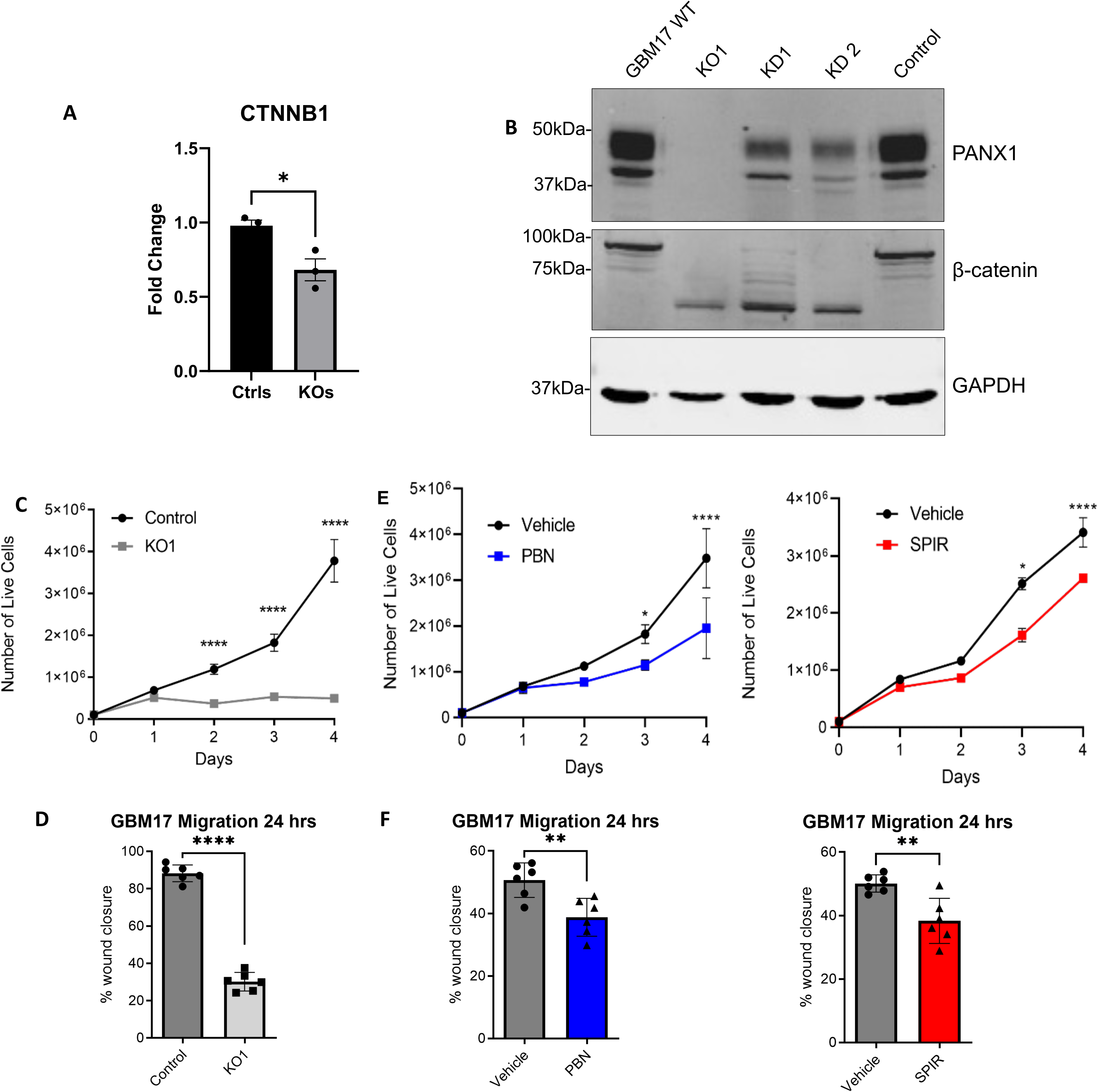
β-catenin protein is disrupted in KO clones and Pannexin 1 drug inhibition reduces cell growth and migration in GBM17 patient-derived cells in 2D. **A)** qPCR showing confirmation of decrease in β-catenin transcript in KO clones vs control clones. **B**) A western blot showing β-catenin protein is disrupted in KO and KD clones compared to control. **C**) Growth curve of a PANX1KO clone compared to control cells shows significantly reduced amount of cells at days 2, 3, and 4 (p<0.0005, N=3, n = 9). **D**) In a scratch wound assay, KO1 clone showed a significantly decreased wound closure compared to control. (N=3, n=9, p>0.0005). **E**) A growth curve of patient-derived cells GBM17 treated with 1 mM Probenecid or 20 µM Spironolactone and respective vehicle controls. Pannexin 1 channel blockers Probenecid (PBN) and Spironolactone (SPIR) both showed a significant decrease in the number of live cells at day 3 (p<0.05) and day 4 (p<0.001). N=3, n=9. **F**) A scratch assay was performed on GBM17 cells and wound closure was measured after 24 hours. PBN and SPIR both reduced the amount of wound closure compared to controls (p<0.05) N=3, n=18.

### Pannexin 1 inhibition changes **β**-catenin localization to a more intracellular pattern

The localization of PANX1 protein appeared to remain the same with both PBN and SPIR blocker treatment, but β-catenin protein shifted from its commonly known membrane location to a more intracellular and perinuclear localization (Fig 6A). However, the total amount of β- catenin protein remained unchanged with PANX1 blocker treatment compared to vehicle controls, indicating a differing effect on the β-catenin protein when there is PANX1 channel inhibition (Fig 6B) versus PANX1 protein deletion (Fig 5B).

**Figure 6.**
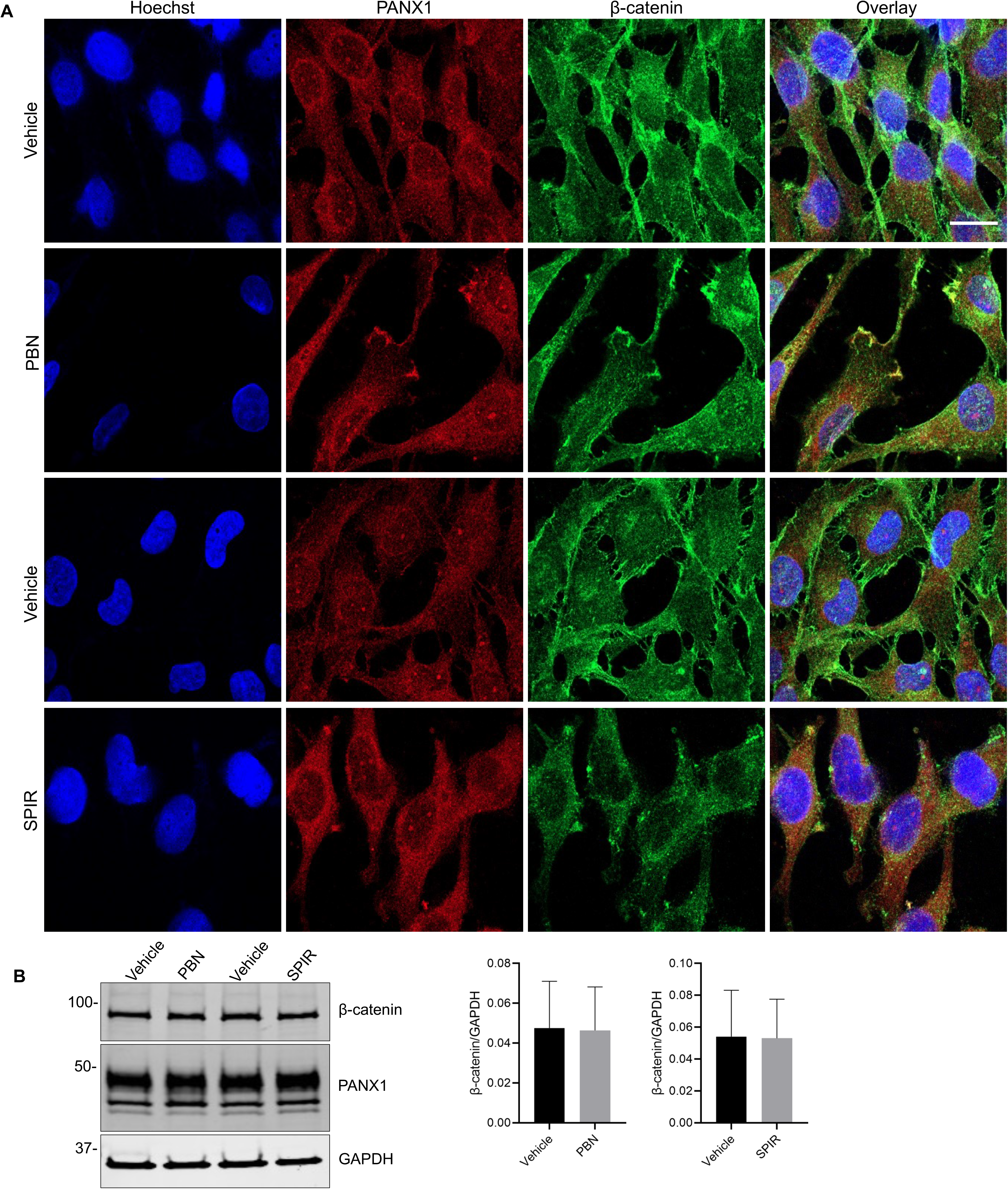
Pannexin 1 inhibition changes β-catenin localization to a more intracellular pattern. **A)** Immunofluorescent staining of patient-derived GBM17 cells treated with 1 mM probenecid, 20 uM spironolactone, or respective vehicle controls over 5 days. GBM17 cells treated with PANX1 inhibitors showed β-catenin localization changing from the cell membrane to more intracellular (green), whereas PANX1 (red) localization was unaffected. Nuclei (blue), N=3, n=9, scale bar = 20 μm. **B**) A western blot of PBN and SPIR treated GBM17 cells showed no difference in PANX1 or β-catenin protein levels compared to respective vehicles (N=3, n = 9).

#### Pannexin 1 inhibition disrupts filamentous actin organization, but not PANX1 localization

Treatment of GBM17 cells with both PBN and SPIR demonstrated a lack of proper F-actin formation visualized by phalloidin staining (Fig 7A). There was no apparent decrease in β-actin protein levels, indicating that the arrangement or polymerization of filamentous actin was disrupted due to other factors (Fig 7B).

**Figure 7.**
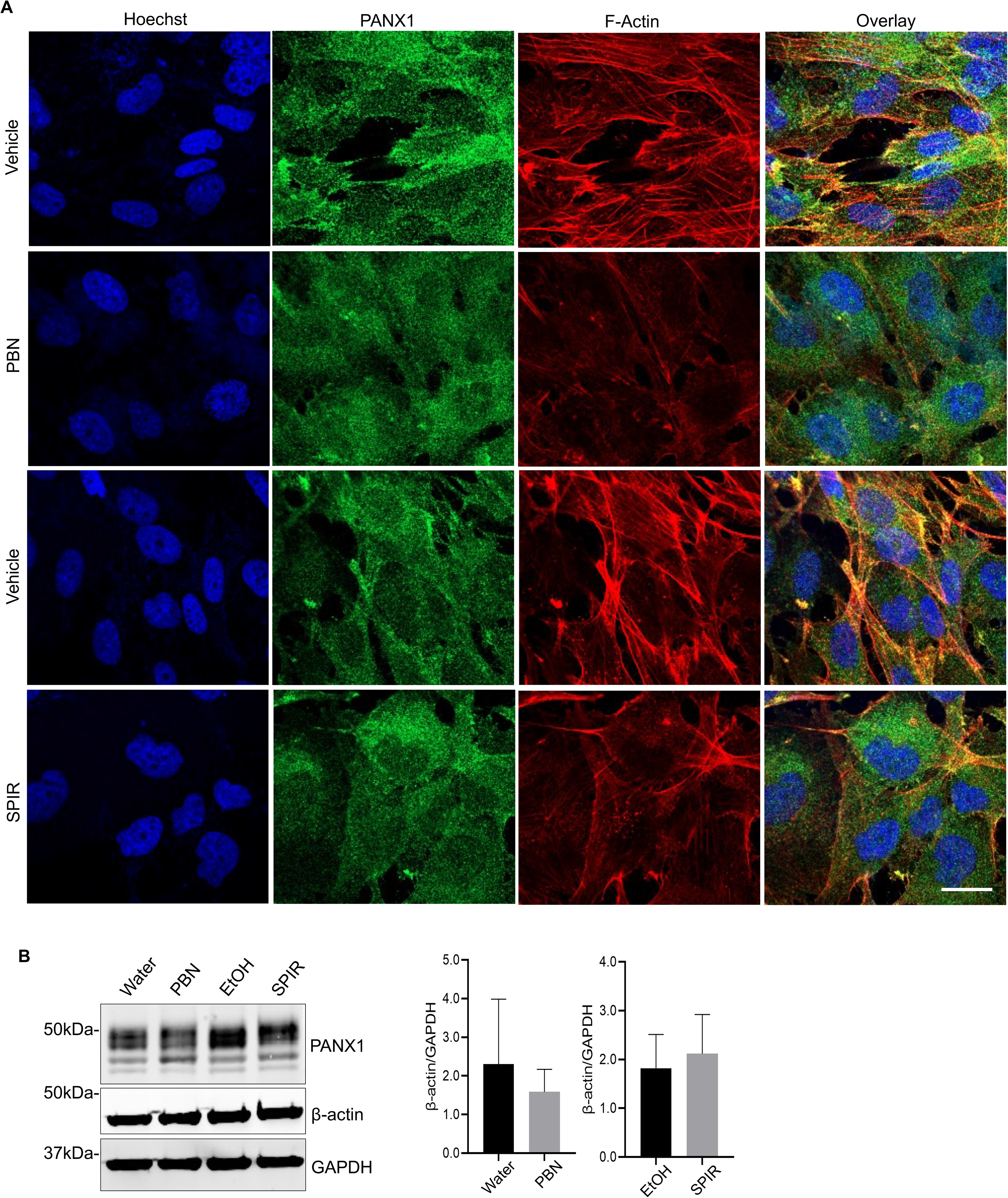
Pannexin 1 inhibition disrupts filamentous actin organization, but not PANX1 localization. **A)** Immunofluorescent staining of patient-derived GBM17 cells treated with 1 mM probenecid, 20 uM spironolactone, or respective vehicle controls over 5 days. GBM17 cells treated with PANX1 inhibitors showed a reduced appearance of F-actin filaments (red), whereas PANX1 (green) localization was unaffected. Nuclei (blue), N=3, n=9, scale bar = 20 μm. **B)** A western blot of PBN and SPIR treated GBM17 cells showed no difference in PANX1 or β- actin protein levels compared to respective vehicles (N=3, n = 9).

### Actin and Pannexin 1 colocalize in invadopodia in GBM cells

LN229 glioblastoma cells were seeded on a gelatin matrix to visualize invadopodia formation. The forming invadopodia could pass through the 1µm pores of the insert (filter) and were visualized by confocal microscopy under the level of cells (Fig 8A-B). As previously published, actin and PANX1 interact directly at PANX1’s C-terminal end. The co-localization of these two proteins was seen in the invadopodia, which may contribute to the invasion and motility of GBM cells (Fig 8C-D).

**Figure 8.**
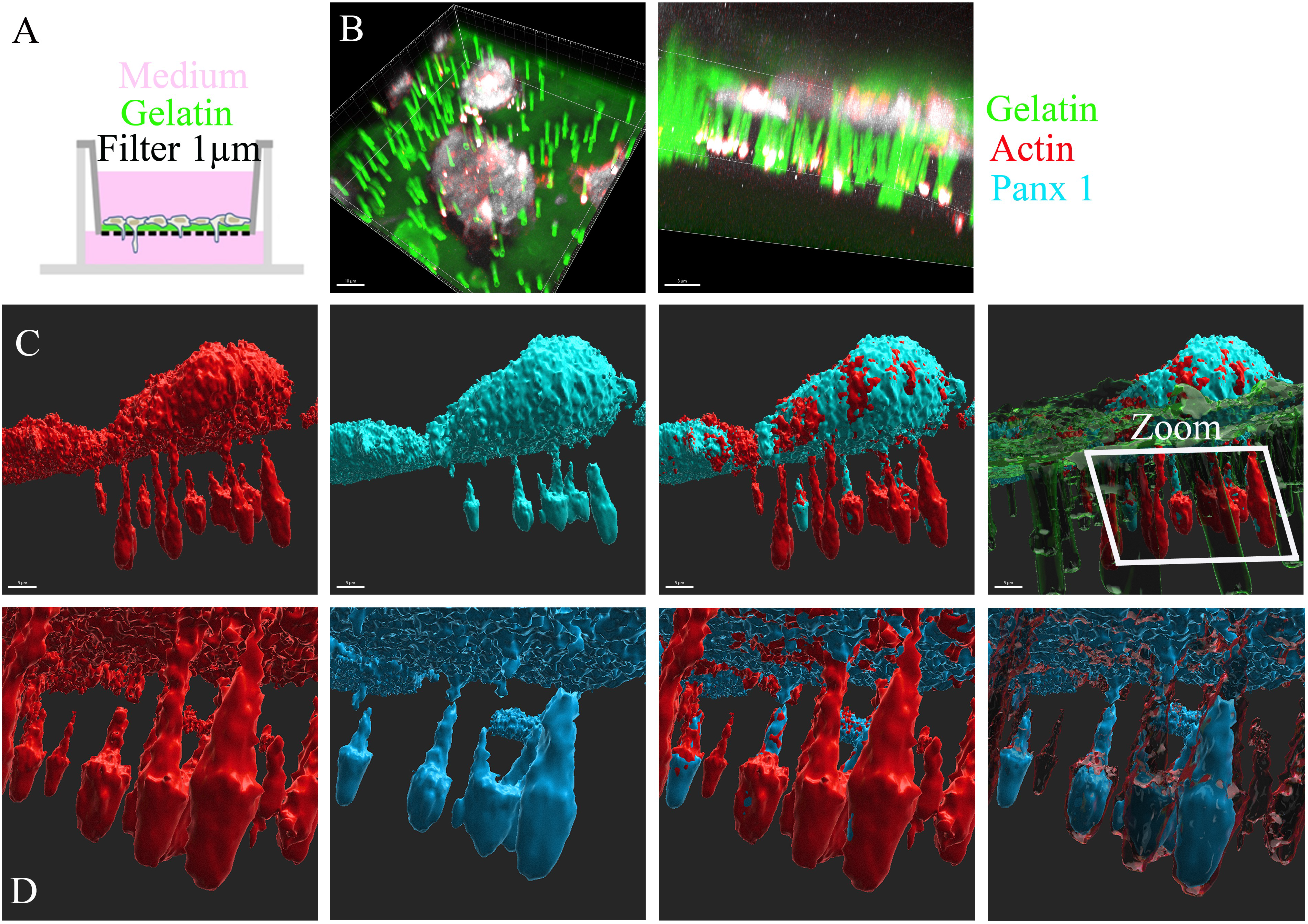
Actin and Pannexin 1 colocalize in invadopodia in GBM cells. **A)** LN229 human glioblastoma cells were seeded on inserts (1 µm-diameter pores) coated with green-Gelatin matrix. **B)** Confocal microscopy images of the Ln229 human glioblastoma cells in *xy* (left panel; scale barre: 10 µm) and *xz* (right panel; scale barre: 8 µm) plans. For a Z dimension, pictures were taken each 400 nm. **C)** Invadopodia were observed below the cells by 3D reconstruction. At their level, Panx1 (blue) and actin (red) were found to be colocalized (Scale Bars: 5 µm). **D)** Different views of the labelled invadopodia (Red: actin; blue: Panx1) from the zoomed square shown in **C** (Right panel).

### PANX1 inhibitors reduce GBM cell growth similar to TMZ treatment but with less cytotoxicity

We wanted to compare PANX1 blocker treatment on GBM cells with the current standard of care TMZ. A growth curve showed PBN reduced cell growth to the same extent as TMZ, while SPIR reduced cell growth compared to control but not to the same degree as TMZ (Fig 9A-B). MTT and Cytotox assays showed a trend towards increased cytotoxicity with TMZ treatment compared to PBN or SPIR inhibition although not significant. This may open avenues to treatments with these drugs in combination (Fig 9C-F).

**Figure 9.**
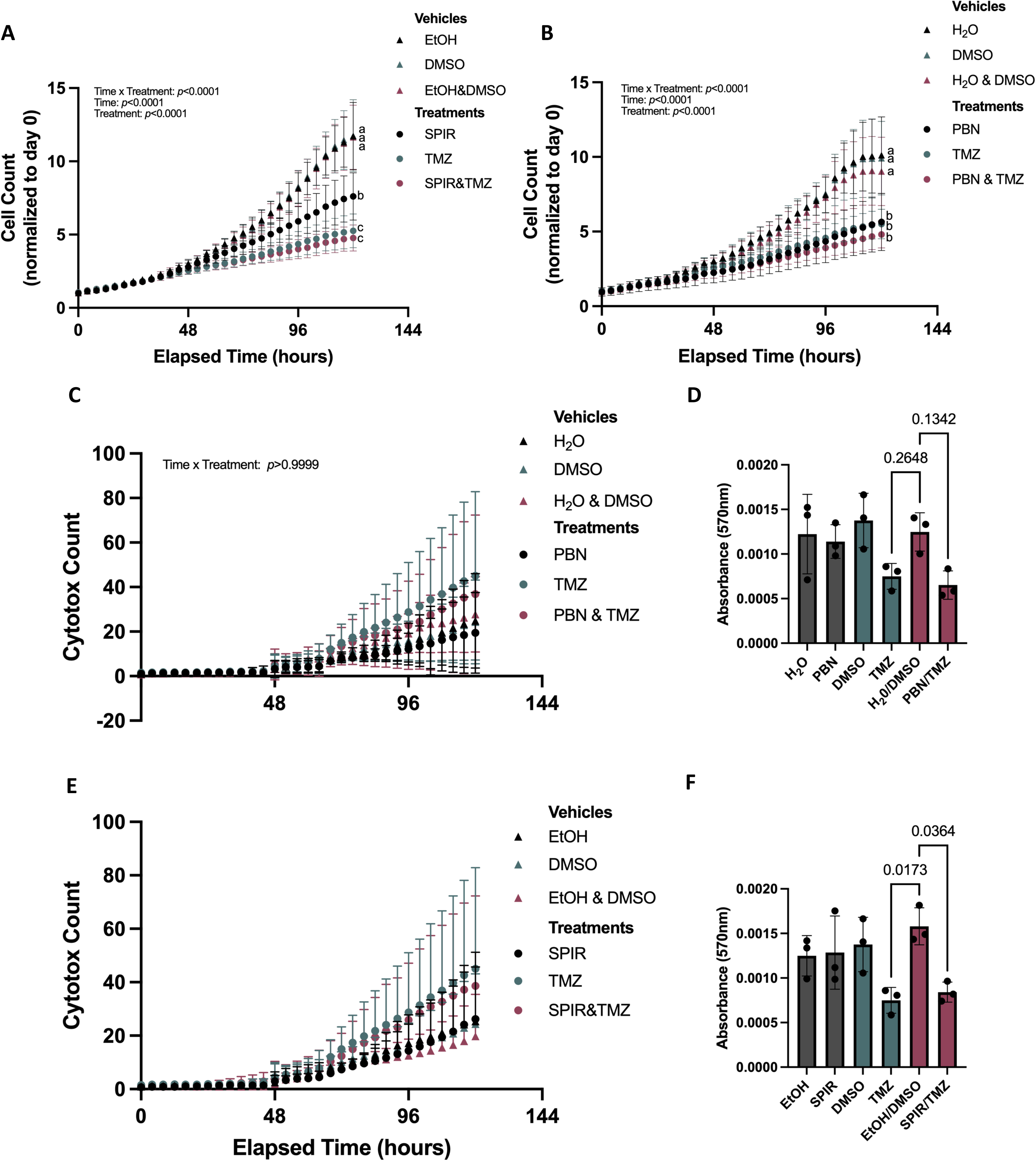
TMZ trends toward exhibiting more toxicity than PANX1 inhibitors. **A)** PBN reduces GBM17 growth to the same extent as TMZ. **B**) SPIR reduces growth to a lesser extent than TMZ but does not alter the effects of TMZ on growth. **C**) MTT assay with GBM17s treated with PBN, TMZ, PBN&TMZ and respective vehicle controls after 5 days. **D**) Cytotox count of GBM17s treated with PBN (1 mM), TMZ (50 µM), PBN&TMZ or vehicle controls (H20, DMSO, H20&DMSO) for five days. No statistically significant differences. N=3, n=18. Count taken on Incucyte S3 System. N=3, n=6. **E**) Cytotox count of GBM17s treated with SPIR (20µM), TMZ (50µM), SPIR&TMZ and vehicle controls (EtOH,DMSO, EtOH&DMSO) over five days. Count taken on Incucyte S3 System. No statistically significant differences. N=3, n=18. **F**) MTT assay with GBM17s treated with SPIR, TMZ, SPIR&TMZ and respective vehicle controls after 5 days. N=3, n=6.

### Blocking Pannexin 1 reduces incidence of tumor hemorrhaging and tumor cell viability in a xenograft chick-CAM model

We used an *ex ovo* chick-CAM model to study the effects of blocking PANX1 channels in a 3D tumor environment. GBM17 cells were mixed with Matrigel and inoculated on a branching vessel of the chick-CAM. After treatment with PBN or vehicle, we observed the control group began hemorrhaging at earlier timepoints compared to PBN (Fig 10A) and had a larger overall number of chicks that hemorrhaged, resulting in nine times greater incidence of hemorrhaging in the control group compared to the PBN treated group (Fig 10B). These tumors were formed with characteristics similar to those in human GBM tumors, as seen with H&E, recapitulating a 3D GBM environment (Fig 10C). New blood vessel formation can promote tumor development by increasing blood supply to the tumor. To explore this result, we excised chick-CAM tumors at endpoint and performed RT-qPCR for key angiogenic markers *VEGFA*, *TGFB1*, *PI3K* and *CXCL8*. We found *IL8* transcripts to be significantly downregulated in the PBN group compared to control (Fig 10D). Lastly, to determine the effect of PANX1 inhibition on tumor viability, GBM17 cells were transduced with LV-tdTomato-Luc2 to express Firefly luciferase. Bioluminescent imaging illustrated a significant decrease in the number of viable cells in the PBN treated group compared to control (Fig 10E).

**Figure 10.**
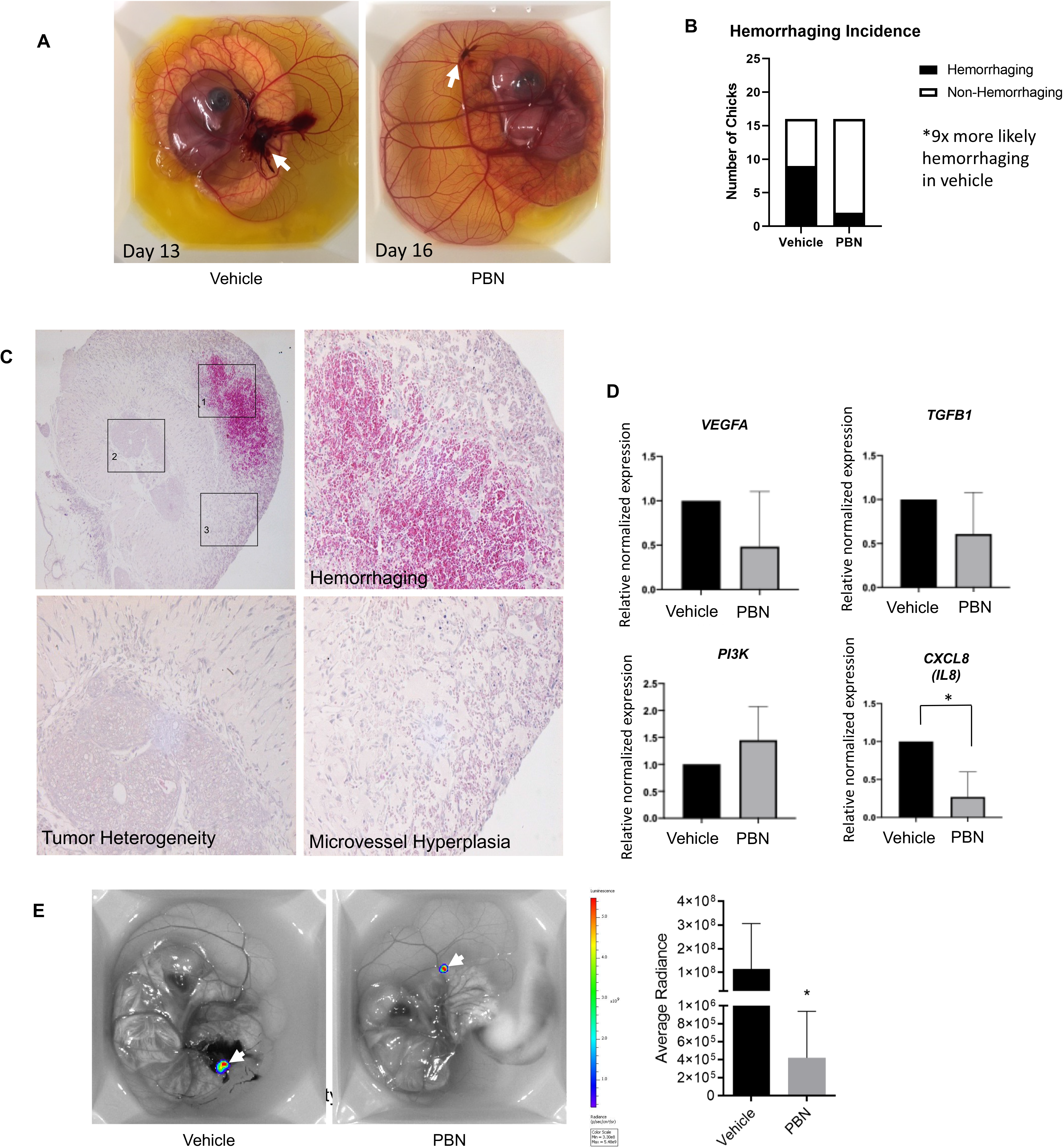
Blocking Pannexin 1 reduces incidence of tumor hemorrhaging and tumor cell viability. GBM17 cells were implanted onto a branching vessel of the chick CAM model on day 10. Tumors were treated with either HBSS or 1mM PBN daily for 7 days. **A)** Tumors treated with PBN visually hemorrhaged less and at a later date compared to vehicle treated tumors. (N=16). **B)** Treatment with PBN resulted in 9x less hemorrhaging incidence compared to the vehicle group. (N=16).**C**) Tumors were excised on day 17 and tissue sections were stained for H&E. Tumor heterogeneity and large necrotic regions with microvessel hyperplasia and coagulation were visualized, which are hallmarks of GBM tumors. **D)** Quantitative PCR for various angiogenic markers on RNA isolated from vehicle and PBN treated chick-CAM tumors. The cytokine IL8 was significantly lower in PBN treated tumors and is involved in vascular leakage. **E)** GBM17 cells were transduced with LV-tdTomato-Luc2 to express Firefly luciferase, and 4 x 10^6^ cells were mixed with Matrigel and implanted onto a branching vessel in the chick CAM model on day 10. Tumors were treated with vehicle (N=12) or 1 mM PBN (N=12) for 7 days, and tumors were evaluated on day 17 by bioluminescent imaging (BLI) on an IVIS Lumina XRMS scanner. A significant reduction in tumor cell viability was observed with PBN treatment.

## DISCUSSION

GBM remains an incurable malignancy with limited therapeutic options, underscoring the need to identify molecular regulators that contribute to tumor growth, invasion, and interactions within the tumor microenvironment [3]. In this study, we extend the evidence supporting PANX1 as a clinically relevant regulator of GBM biology by integrating patient tumor observations with genetic, pharmacologic, and *ex vivo* xenograft approaches. Across these systems, PANX1 perturbation consistently reduced GBM cell fitness, altered cytoskeletal organization, and modified tumor-associated vascular phenotypes. Rather than acting solely as a generic proliferative driver, our data support a model in which PANX1 helps maintain a tumor-promoting “interface state” characterized by coordinated regulation of cell-cell and cell-ECM interactions, β-catenin organization, actin architecture, and inflammatory/vascular crosstalk. This interpretation is supported by the convergence of phenotypes observed after CRISPR/Cas9- mediated PANX1 deletion and pharmacologic inhibition with two repurposed agents, PBN and SPIR.

The role of PANX1 in glioma has remained somewhat context-dependent across the literature. In rat C6 glioma cells, restoration of PANX1 was previously reported to suppress proliferation, motility, and anchorage-independent growth, suggesting a tumor-suppressive role in that engineered setting [23]. However, subsequent work showed that PANX1 can promote ATP-dependent actomyosin remodeling and accelerate multicellular aggregate compaction through purinergic signalling, highlighting its ability to function as a biomechanical organizer rather than a simple binary suppressor or promoter [24]. In more human-relevant glioma systems, PANX1 silencing in U87-MG cells reduced proliferation and decreased inflammatory mediator output, including IL-6, IL-8, and glutamate release [22, 45]. Additional studies have also implicated PANX1 in pro-tumorigenic regulatory axes in glioma, including circRNA/miRNA-linked pathways [25]. Our findings in patient-derived GBM models align more closely with this latter body of work and strengthen the interpretation that, in human GBM contexts, PANX1 supports pro-tumor-associated behaviours. At the same time, the apparent discrepancy with the C6 literature likely reflects species differences between rat and human, baseline expression state, and the biological consequence of the perturbation used.

A major mechanistic theme emerging from our data is that PANX1 loss disproportionately disrupts programs linked to the cell surface, adhesion, and ECM signalling interface. Transcriptomic analyses of PANX1-KO cells identified enrichment changes in pathways such as ECM-receptor interaction, focal adhesion, and adhesion-related gene ontology categories, consistent with a role for PANX1 in maintaining signalling competence at the plasma membrane. In GBM, these pathways are not merely structural; they are tightly coupled to invasive behaviour, cell survival, and therapy resistance through focal adhesion kinase, PI3K/AKT, and related signalling nodes [46]. Accordingly, the reduced growth and impaired scratch wound closure we observed in PANX1-deficient cells can be interpreted as coordinated consequences of destabilizing this interface state rather than isolated effects on proliferation alone. This view also fits with the known biology of PANX1 as a channel capable of shaping local purinergic microenvironments via ATP release [10], thereby influencing paracrine and autocrine signalling at sites of cell-cell and cell-ECM communication [6].

Our data further support a mechanistic link between PANX1, β-catenin organization, and actin dynamics. Genetic loss of PANX1 disrupted β-catenin at both the transcript and protein levels, whereas pharmacologic inhibition altered β-catenin localization and reduced filamentous actin organization without major changes in total β-actin abundance. This convergence strongly suggests that PANX1 contributes to maintenance of a cytoskeletal and adhesion-permissive state. Prior work provides a plausible physical basis for this interpretation: PANX1 has been shown to interact with actin microfilaments, and intact actin architecture is important for PANX1 localization and mobility at the cell surface [17, 47]. PANX1 has also been linked to Arp3- containing cytoskeletal complexes, supporting a bidirectional relationship in which cytoskeletal integrity stabilizes PANX1 while PANX1 function feeds back to influence actin remodelling [47]. Our observation that PANX1 localizes with actin-rich invadopodia in LN229 cells extends this concept to invasive GBM structures and suggests that PANX1 may be enriched at sites where local matrix interaction, protrusive actin polymerization, and signalling are most tightly integrated.

The β-catenin findings are particularly notable given the established role of Wnt/β- catenin signalling in GBM proliferation, stemness, invasion, and therapeutic resistance [48, 49]. Our results are consistent with a model in which PANX1 supports β-catenin stability, trafficking, or membrane-associated organization, thereby helping maintain downstream transcriptional and adhesive programs. A useful precedent comes from melanoma, where PANX1 was shown to engage β-catenin and regulate tumor cell growth and metabolic activity [26]. Although GBM and melanoma are biologically distinct, this finding supports the broader idea that PANX1 can regulate oncogenic phenotypes through β-catenin-associated mechanisms. The distinction we observed between knockout and inhibitor-treated conditions may also be biologically informative. Complete PANX1 deletion likely removes both channel-dependent and channel-independent functions and permits long-term network adaptation, whereas drug treatment may only partially inhibit channel activity over shorter timescales. Thus, the intracellular redistribution of β-catenin seen with PBN or SPIR may reflect altered adhesion dynamics and cytoskeletal tension, while the more profound β-catenin change seen in KO cells may require chronic loss of PANX1-dependent scaffolding and signalling functions.

The Hippo pathway provides an additional mechanistic bridge linking these cytoskeletal effects to broader transcriptional rewiring as we have reported in melanoma [50]. YAP/TAZ are well-established sensors of actomyosin tension and ECM mechanics, and their activity is closely linked to invasive and pro-growth states in GBM [51, 52]. The downregulation of Wnt/β-catenin- and YAP/TAZ-associated signatures in our PANX1 knockout transcriptomic data is therefore consistent with PANX1 supporting a mechanically permissive state that enables pro-invasive transcriptional outputs. In this framework, PANX1 may act at two interconnected levels: first, by contributing directly to cytoskeletal organization through protein interactions and ATP-dependent purinergic signaling; and second, by helping sustain adhesion-associated signalling hubs that reinforce β-catenin and YAP/TAZ activity. This model fits well with prior evidence that PANX1 can remodel actin through ATP-P2X_7_ signalling [24] and provides a plausible explanation for why PANX1 perturbation simultaneously affects growth, migration, and interface-associated transcriptional programs.

An important translational implication of this study is that pharmacologic PANX1 inhibition reproduced key, but not all, aspects of the genetic phenotype. Both PBN and SPIR reduced growth and migration and altered β-catenin/actin organization, supporting the idea that channel blockade can phenocopy core elements of PANX1 loss. This convergence is meaningful because PBN has long been used as a functional PANX1 inhibitor [27], while SPIR was more recently identified as a potent PANX1 blocker independent of its canonical mineralocorticoid receptor activity [28]. At the same time, neither drug is fully PANX1-exclusive, and PBN in particular, can alter other cellular processes, including PANX1-associated protein interactions [53]. For this reason, our pharmacologic data are best interpreted as PANX1-enriched rather than PANX1-specific. Notably, the incomplete overlap between genetic and pharmacologic phenotypes may even reinforce the likelihood that PANX1 contributes through both channel-dependent and channel-independent mechanisms.

The comparison with TMZ is also informative. TMZ tended toward greater acute toxicity than either PANX1 inhibitor, and combination treatment did not produce a strong additive effect in the endpoints examined. Rather than weakening the therapeutic rationale, this may indicate that PANX1 inhibition primarily acts by suppressing invasion-associated, cytoskeletal, and microenvironmental programs rather than by serving as a strongly cytotoxic monotherapy in conventional 2D assays [54]. This distinction may be especially relevant in GBM, where standard therapy can enrich for more invasive post-treatment survivors and promote invadopodia-associated behaviours [52, 55]. In that context, PANX1-directed therapy may be most valuable as an invasion-modulating adjunct rather than as a direct cytotoxic substitute.

One of the most distinctive findings of this study is the reduction in hemorrhagic/leak phenotypes in the chick-CAM model following PBN treatment, accompanied by reduced tumor cell viability and decreased IL-8/CXCL8 expression. This observation extends the role of PANX1 beyond tumor-intrinsic growth control and suggests that PANX1 may also influence tumor-vascular interactions, which are areas of PANX1 localization (Fig 1C). GBM vasculature is highly abnormal and prone to permeability, edema, and hemorrhagic instability, making the reduction in CAM hemorrhaging biologically meaningful. IL-8 is a plausible mechanistic mediator here, as GBM-derived IL-8 has been shown to increase endothelial permeability and disrupt junctional integrity [56, 57]. The reduction in IL-8 expression in treated chick-CAM tumors is also directionally consistent with prior glioma studies reporting decreased IL-8 after PANX1 silencing [45]. Furthermore, RNA-seq enrichment of inflammatory response signatures in PANX1-KO GBM cells indicate that PANX1 activity modulates this response (Fig S3). Thus, our chick-CAM data support a model in which PANX1 contributes not only to GBM cell fitness and invasion, but also to inflammatory outputs that may exacerbate vascular leakiness in the tumor microenvironment.

Taken together, our data support a model in which PANX1 functions as a nexus linking GBM growth, cytoskeletal organization, invasive interface biology, and vascular/inflammatory phenotypes. By integrating patient-derived genetic perturbation, convergent pharmacologic inhibition, invadopodia-associated imaging, and chick-CAM vascular phenotyping, this study moves PANX1 beyond a purely descriptive channel biology framework and positions it as a mechanistically relevant regulator of GBM cell state and tumor-microenvironment interaction. Future separation-of-function mutants, direct readouts of β-catenin, YAP/TAZ, and endothelial permeability will be important to refine causality. Nevertheless, the current dataset already supports PANX1 as a promising translational node whose disruption may impair GBM progression by destabilizing the adhesive cytoskeletal, and inflammatory programs that sustain aggressive disease.

## Data availability statement

All data is included in the current manuscript, and transcriptomic data is available from the NCBI GEO (Gene Expression Omnibus) repository, Accession # GSE337204.

## Acknowledgements and Funding

We thank the funding agencies that made this work possible including the following: Canadian Institutes of Health Research (CIHR; PJT 185953 to SP, MH, and JR). Catalyst grant from London Cancer Research Program (LRCP) to SP, MH and JR, Brain Tumour Foundation grant R5313A22 to SP. Ligue contre le Cancer, Comité de la Vienne to ND and MM.

## Author contributions

DJ: study design, data acquisition, analysis, manuscript writing and editing

MH: Data acquisition, analysis, manuscript writing and editing

RK, CVK, JK, RSP, BO’D, RL, CH: data generation, analysis, methodology, manuscript editing

ND: Data acquisition, data analysis, manuscript editing

AD, MM, JR: supervision, data analysis, manuscript editing

MH: Acquisition and culture of patient tissue specimens, manuscript editing

SP: Study design, funding acquisition, supervision, data acquisition, analysis, manuscript editing

## Supplementary Figures

**S1. Wnt.** β**-catenin, F-actin, YAP/TAZ are downregulated with Pannexin 1 knock out.** Differential gene expression within the hippo signalling pathway in PANX1-KO compared to control GBM cells. Upregulated (fold change ≥ 1.5) genes are represented in red and downregulated genes are represented in blue (fold change ≤ -1.5). Differentially expressed genes have a p-value ≤ 0.05.

**S2. qPCR confirmation of selected top DEGs A)** Volcano plot showing DEGs – Top 12 DEGs labelled. **B)** Principal component analysis (PCA) plot indicating two distinct clusters for PANX1-KO and control GBM clones. **C)** MA plot with LFC shrinkage visualizing the differential expression of detected genes (P≤0.05 [blue], non- significant [grey]). Log(Fold Change) values beyond the axis boundaries are indicated by a triangle.**D)** qPCR on RNA from KO and Cntrl clones was performed to confirm some top DEGs from Partek analysis.

**S3. PANX1 knockout in primary GBM cells is associated with proliferation, apoptosis, and inflammatory response pathways.** Significantly enriched hallmark gene sets determined by Gene Set Enrichment Analysis (GSEA) of normalized counts output from DESeq2. Enrichment plots provided for the 9 gene sets significant at FDR < 25%. Nominal p-values are listed above each enrichment plot, calculated by GSEA, highlighting eight gene sets significantly enriched in the PANX1-KO phenotype at nominal p-value < 0.05, out of 50 defined hallmark gene sets.

**S4. Top 50 DEGs in PANX1-KO versus control GBM clones, ranked by FDR step-up.** Table of top 50 DEGs. Data collected using RNA-seq and analyzed using DESeq2 in Partek Flow. Description of genes generated with g:Profiler. Ensembl ID is provided in place of Gene ID for lncRNAs.

