## Supplementary material for "Pannexin 1 inhibition reduces tumorigenic properties of patient-derived glioblastoma cells through the HIPPO and Wnt signalling pathways": Suppl figures

### HIPPO SIGNALING PATHWAY

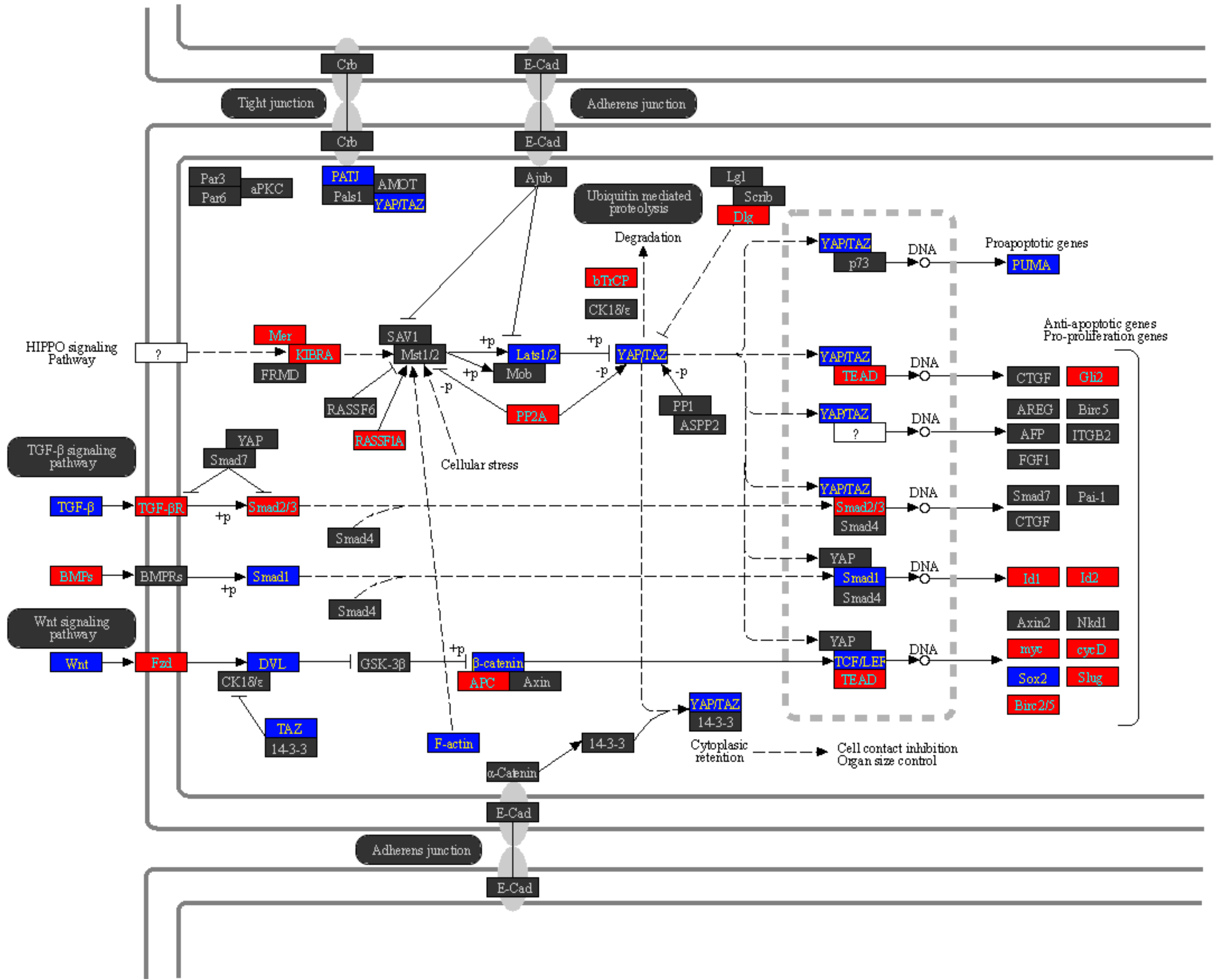

A

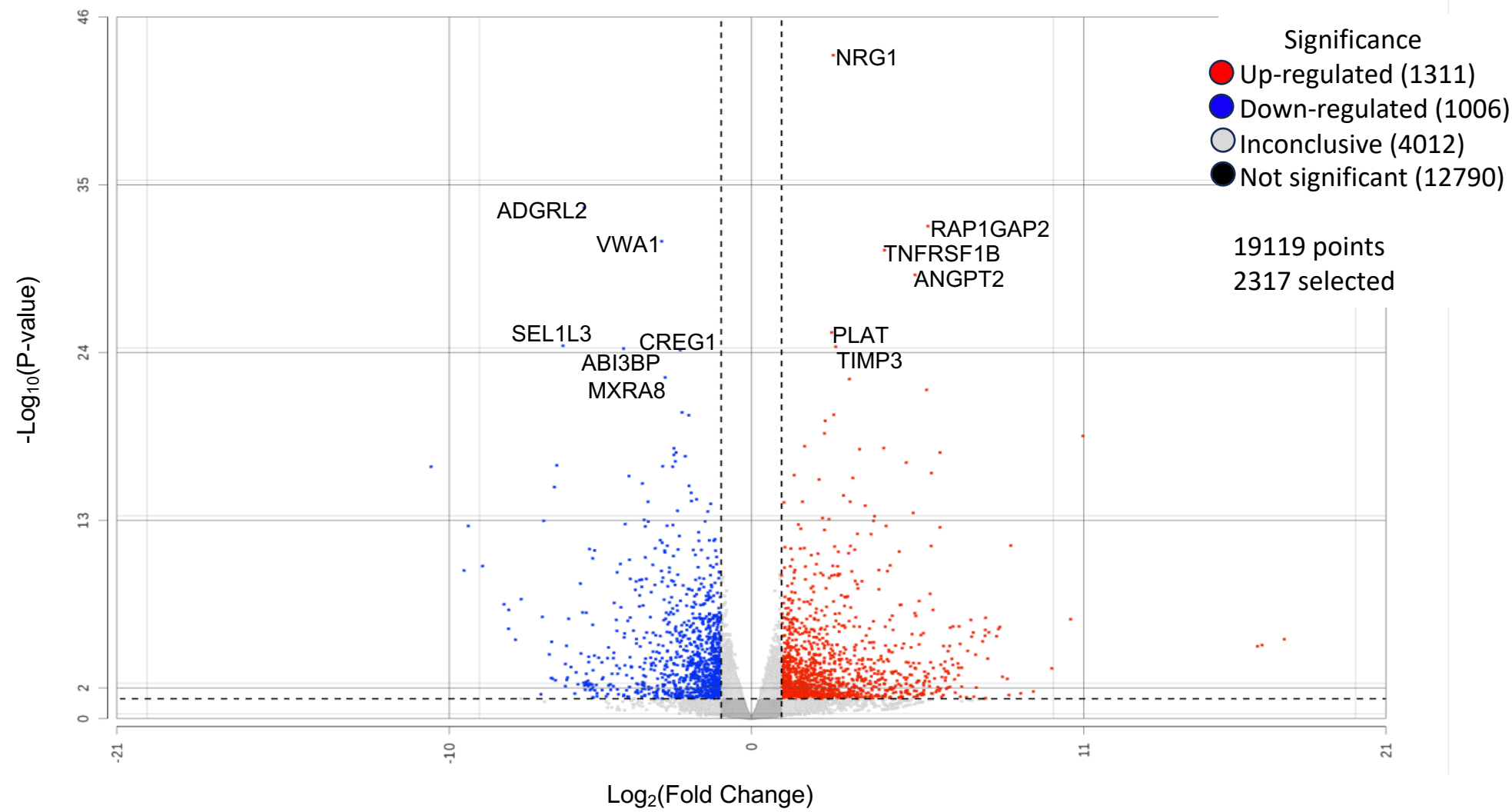

B

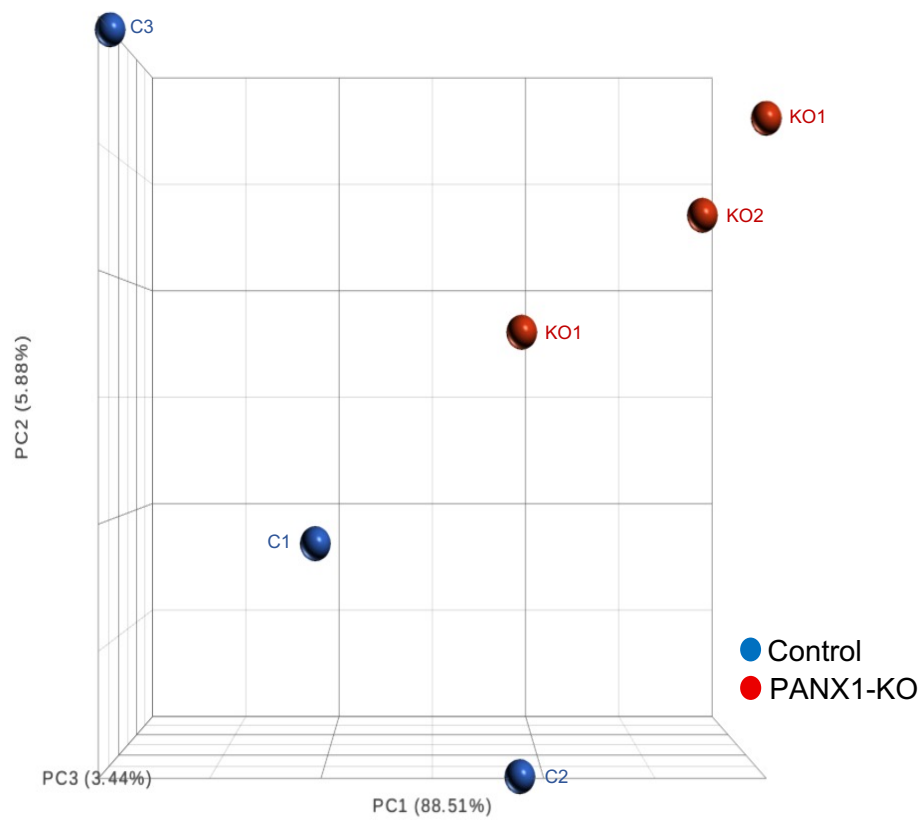

C

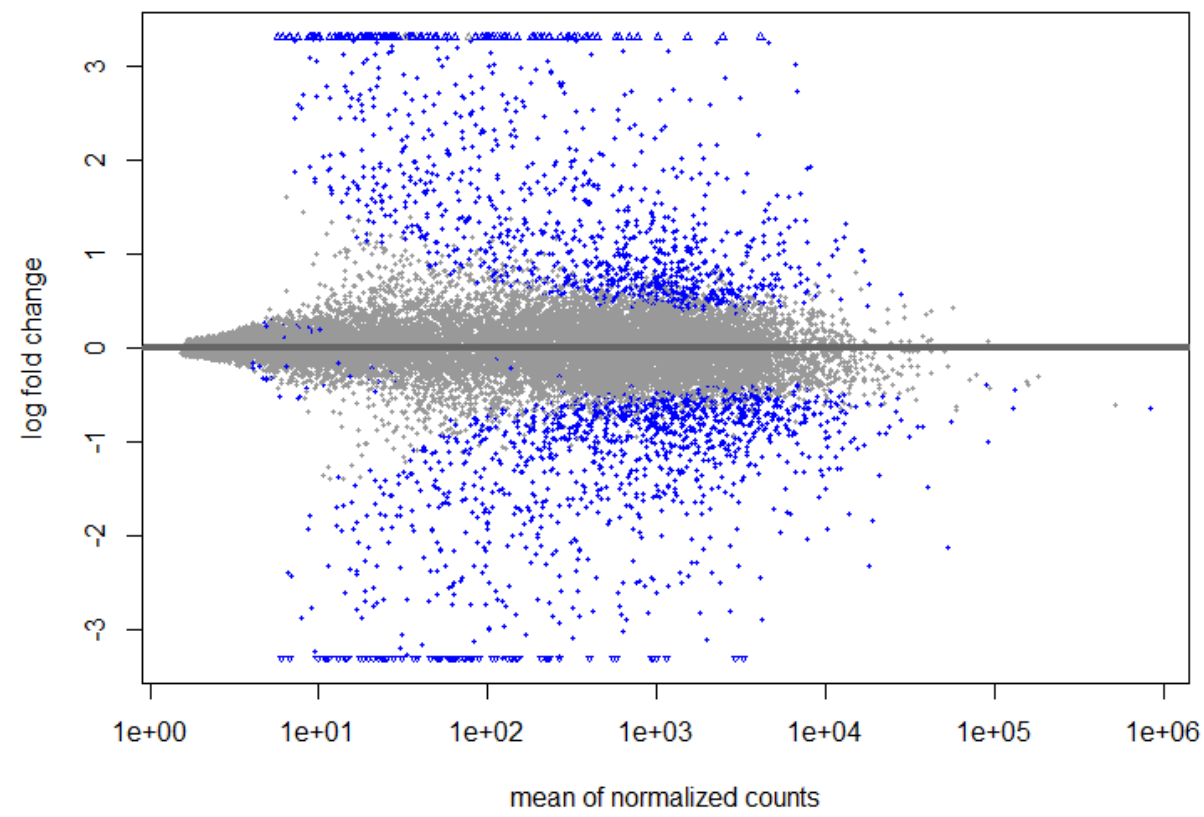

D

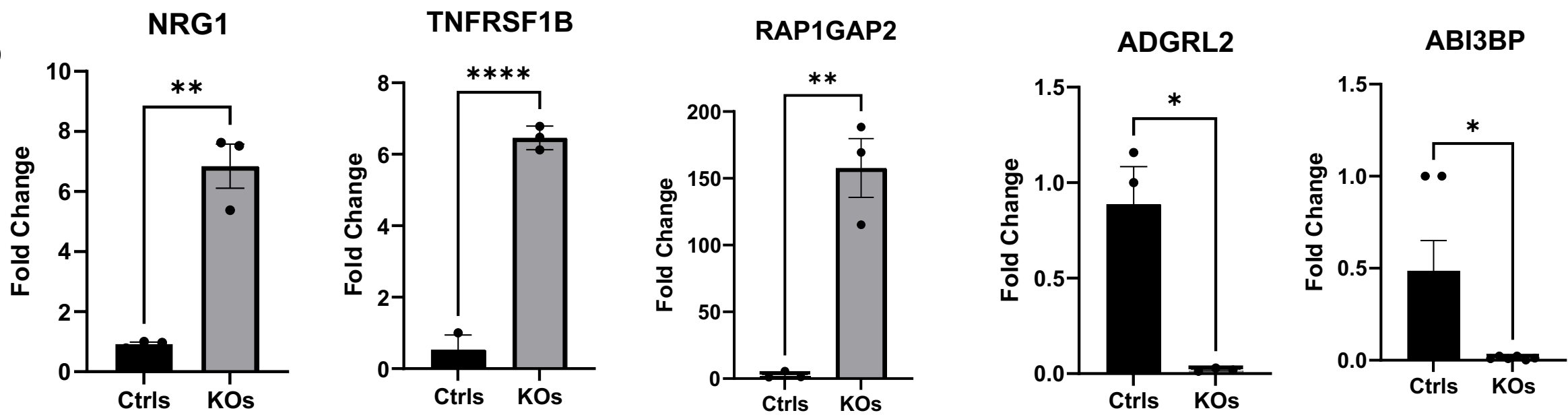

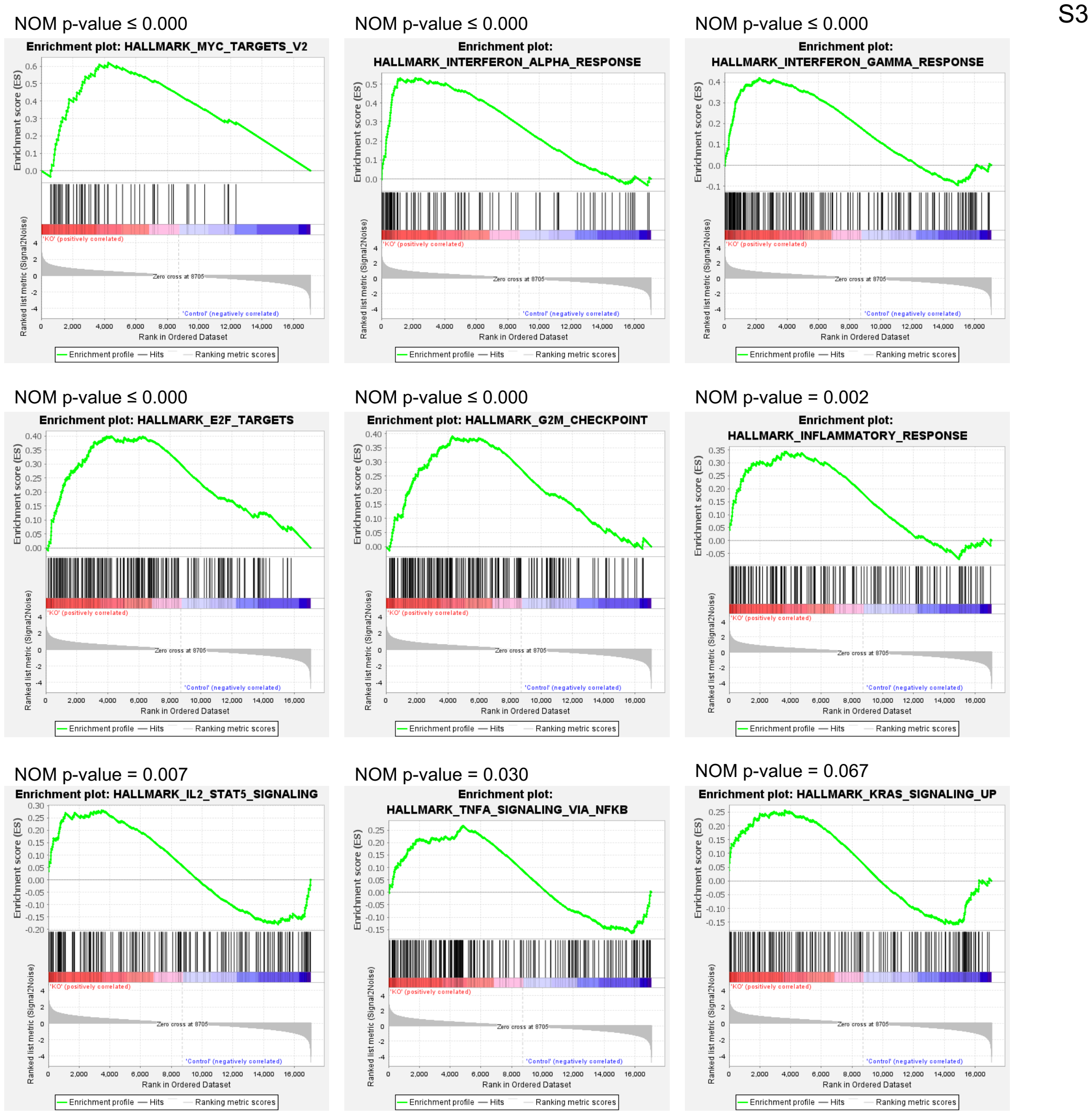

| Gene Name | Description | FDR step-up | Log<sub>2</sub>(Ratio) |
| --- | --- | --- | --- |
| NRG1 | Neuregulin 1 | 5.84E-40 | 2.71 |
| ADGRL2 | Adhesion G protein-coupled receptor L2 | 2.97E-30 | -5.54 |
| RAP1GAP2 | RAP1 GTPase activating protein 2 | 3.17E-29 | 5.84 |
| VWA1 | von Willebrand factor A domain containing 1 | 2.36E-28 | -2.97 |
| TNFRSF1B | TNF receptor superfamily member 1B | 7.06E-28 | 4.40 |
| ANGPT2 | Angiopoietin 2 | 2.44E-26 | 5.42 |
| PLAT | Plasminogen activator, tissue type | 1.27E-22 | 2.65 |
| SEL1L3 | SEL1L family member 3 | 8.18E-22 | -6.24 |
| TIMP3 | TIMP metalloproteinase inhibitor 3 | 8.54E-22 | 2.79 |
| ABI3BP | ABI family member 3 binding protein | 1.02E-21 | -4.23 |
| CREG1 | Cellular repressor of E1A stimulated genes 1 | 1.16E-21 | -2.36 |
| MXRA8 | Matrix remodeling associated 8 | 6.60E-20 | -2.85 |
| PAQR5 | Progestin and adipoQ receptor family member 5 | 7.87E-20 | 3.24 |
| RSAD2 | Radical S-adenosyl methionine domain containing 2 | 3.67E-19 | 5.80 |
| LFNG | LFNG O-fucosylpeptide 3-beta-N-acetylglucosaminyltransferase | 1.03E-17 | -2.30 |
| RIGI | RNA sensor RIG-I | 1.38E-17 | 2.72 |
| HMGCS1 | 3-hydroxy-3-methylglutaryl-CoA synthase 1 | 1.41E-17 | -2.07 |
| LRRC8C | Leucine rich repeat containing 8 VRAC subunit C | 3.05E-17 | 2.44 |
| CNNM1 | Cyclin and CBS domain divalent metal cation transport mediator 1 | 1.96E-16 | 2.42 |
| SOX11 | SRY-box transcription factor 11 | 2.72E-16 | 10.97 |
| LINC-PINT | Long intergenic non-protein coding RNA, p53 induced transcript | 1.24E-15 | 1.76 |
| PPM1E | Protein phosphatase, Mg<sup>2+</sup>/Mn<sup>2+</sup> dependent 1E | 1.52E-15 | 4.38 |
| MTUS1 | Microtubule associated scaffold protein 1 | 1.52E-15 | -2.56 |
| SGIP1 | SH3GL interacting endocytic adaptor 1 | 1.69E-15 | 3.58 |
| MAGED4 | MAGE family member D4 | 2.54E-15 | 6.24 |
| NDRG1 | N-myc downstream regulated 1 | 2.54E-15 | -2.49 |
| DDIT4 | DNA damage inducible transcript 4 | 3.51E-15 | -2.56 |
| PRUNE2 | Prune homolog 2 with BCH domain | 4.15E-15 | -2.19 |
| NREP | Neuronal regeneration related protein | 8.72E-15 | -2.52 |
| RIPOR2 | RHO family interacting cell polarization regulator 2 | 9.93E-15 | 5.12 |
| WNT9A | Wnt family member 9A | 1.47E-14 | -6.44 |
| SSC5D | Scavenger receptor cysteine rich family member with 5 domains | 1.66E-14 | -2.94 |
| F11R | F11 receptor | 1.66E-14 | -2.61 |
| PCDH20 | Protocadherin 20 | 1.66E-14 | -10.60 |
| RELN | Reelin | 4.16E-14 | 5.96 |
| NPC1 | NPC intracellular cholesterol transporter 1 | 5.64E-14 | 1.41 |
| ACTBL2 | Actin beta like 2 | 6.37E-14 | -4.05 |
| HGF | Hepatocyte growth factor | 8.10E-14 | 3.36 |
| HELZ2 | Helicase with zinc finger 2 | 9.92E-14 | 2.24 |
| CSF1R | Colony stimulating factor 1 receptor | 1.78E-13 | -3.61 |
| PLEKHO2 | Pleckstrin homology domain containing O2 | 2.51E-13 | -2.06 |
| EDNRB | Endothelin receptor type B | 2.94E-13 | -6.52 |
| ENSG00000280800 | Novel transcript, lncRNA | 6.70E-13 | -1.99 |
| STMN3 | Stathmin 3 | 9.90E-13 | 3.05 |
| ENSG00000280614 | Novel transcript, lncRNA | 1.74E-12 | -1.81 |
| ENSG00000281181 | Novel transcript, lncRNA | 2.17E-12 | -1.98 |
| SAMD9 | Sterile alpha motif domain containing 9 | 2.31E-12 | 1.69 |
| ISG15 | ISG15 ubiquitin like modifier | 2.31E-12 | 3.27 |
| COL4A1 | Collagen type IV alpha 1 chain | 2.31E-12 | -3.42 |
| PRKDC | Protein kinase, DNA-activated, catalytic subunit | 2.46E-12 | 1.08 |
